# 3-Hydroxypropanamidines: from antiplasmodial leads to a novel antibacterial scaffold

**DOI:** 10.64898/2026.09.20.752996

**Authors:** Lara-Sophie Kaufmann, André Schultz, Noemi Wladarz, Tanja Gangnus, Beate Lungerich, Tanja C. Knaab, André Haase, Lea Joswig, Finn Dressler, Pascal Dietze, Ralf Benndorf, Bjoern B. Burckhardt, Thomas Kurz, Julia E. Bandow

## Abstract

Arylamino alcohols constitute an established class of antimalarial agents. While this class is best known for its potent antiplasmodial activity, selected members, most notably mefloquine, have also been reported to exhibit antibacterial activity. Structurally related 3-hydroxypropanamidines were developed as highly potent antimalarial agents. Here, we explored the antibacterial activity of three 3-hydroxypropanamidines (**4** (**BLK278**), **5** (**BLK280**), and **6** (**TKK088**)) against clinically relevant Gram-positive and Gram-negative bacteria and investigated their cytotoxicity, *in vitro* pharmacokinetic properties, and proteome-based bacterial response to treatment. We observed good antibacterial activity against *Staphylococcus aureus*, with **6** (**TKK088**) displaying the highest activity against multidrug-resistant MRSA and VISA strains. **5** (**BLK280**) demonstrated the greatest spectrum of activity acting also against clinically relevant Gram-negative strains of *Escherichia coli*, *Acinetobacter baumannii*, *Klebsiella pneumoniae*, *Klebsiella aerogenes*, and *Shigella sonnei*. In cytotoxicity assays with HUVEC and HAOSMC cells, **4** (**BLK278**) and the previously reported **7** (**TKK129**) showed the most favourable overall profiles among the investigated 3-hydroxypropanamidines. The selectivity indices for *S. aureus* were below 2.5 with clear cell-type-dependent differences for all compounds including mefloquine, suggesting that improving antibacterial selectivity is an important objective for further optimization. *In vitro* pharmacokinetics showed high plasma protein binding and good human microsomal and plasma stability for all three 3-hydroxypropanamidines. Gel-based proteomic responses revealed a strong impact on the bacterial cell envelope. The observed acute proteomic responses reflect pleiotropic effects and are consistent with earlier reports implicating F_0_F_1_-ATPase as a potential molecular target of mefloquine in *Streptococcus pneumoniae*.

## Introduction

The WHO repeatedly pointed out antimicrobial resistance as a serious global health threat [1]. Yet, very few new antibiotic classes have been introduced to the market in the last decades [2]. This has led to studies exploring the possibility of repurposing known and approved therapeutics from different medical indications as antibiotics. In one such screening effort by Kunin and Ellis, mefloquine, an arylamino alcohol, was found to be active against *Streptococcus pneumoniae* [3]. Mefloquine was originally used for treatment and prophylaxis against malaria, which is caused by the eukaryotic parasite *Plasmodium falciparum*. The arylamino alcohols quinine, mefloquine, and lumefantrine inhibit hemozoin formation [4]. Mefloquine was further shown to inhibit the 80S ribosome of *P. falciparum* by interacting with the GTPase-associated centre, binding to rRNA, protein, and the coordinating Mg^2+^ [5]. Among clinically used arylamino alcohol antimalarials mefloquine stands out because of its documented antibacterial activity, whereas comparable evidence for quinine, halofantrine, and lumefantrine is weak or lacking. The potential application of mefloquine as antibacterial drug has been limited by concerns about neuropsychiatric adverse effects [6–8]. Nevertheless, these studies laid the groundwork showing that antimalarial agents could have antibacterial activity. In *S. pneumoniae*, mutation mapping identified F_0_F_1_-ATPase as potential molecular target of mefloquine [9]. In *Mycobacterium abscessus*, *in silico* docking studies predicted a strong interaction with a resistance-nodulation-division (RND) protein, the mycolic acid transporter MmpL3 [10], and in cell-based assays *Escherichia coli* and *Pseudomonas aeruginosa* RND efflux pump activity was found to be diminished, and pleiotropic effects on energy metabolism and biofilms as well as synergistic effects with antibiotics like colistin were observed [11–13]. To the best of our knowledge, experimental evidence of interaction of mefloquine or any other arylamino alcohol with a bacterial target is still lacking. However, many of the reported effects could be explained by interference with utilization of the proton motive force and/or ATP synthesis by bacterial F_0_F_1_-ATPase. Mammalian cells have a homologous F_0_F_1_-ATPase in mitochondria, the structure of which differs from that of bacteria in the reportedly relevant membrane-integrated F_0_ subunit. The mitochondrial F_0_F_1_-ATPase has a *c*_8_ ring [14], that of *E. coli* a *c*_10_ ring [15] and *Bacillus c*-ring sizes differ by species, with *c*_10_ and *c*_13_-rings reported for *Bacillus* PS3 and *Bacillus sp.* strain TA2.A1, respectively [16].

Like mefloquine, lumefantrine and halofantrine belong to the chemical class of arylamino alcohols [17]. Like mefloquine, halofantrine is only of limited clinical relevance due to adverse cardiac events [18]. Lumefantrine on the other hand remains of high clinical relevance. The 3-hydroxypropanamidine (3-HPA) scaffold originated from the medicinal chemistry optimization of the previously developed 3-hydroxypropanehydrazonamide series, which emerged from a phenotypic antimalarial screening campaign at the Bernhard Nocht Institute for Tropical Medicine (BNITM, Hamburg)[19, 20]. Continued lead optimization of this scaffold ultimately yielded the preclinical antimalarial lead compounds **7** (**TKK129**) and TKK130, which demonstrated excellent oral efficacy and tolerability in murine malaria models [21, 22]. Both showed excellent antimalarial activity *in vitro* (in the low nanomolar range), low cytotoxicity against immortalized human cell lines (HepG2, HeLa, HEK293), good pharmacokinetics in terms of metabolic stability, plasma-protein binding, and snapshot *in vivo* pharmacokinetics in mice. Most notably, in mouse models of infection, **7 (TKK 129)** and TKK130 showed excellent *in vivo* efficacy in a standard 4-day Peters test in *Plasmodium berghei*-infected mice with once daily oral dosing at 3 mg/kg up to 50 mg/kg for four consecutive days [21, 22]. Reduction in parasite loads determined on day four and safety at day 30 were excellent with no obvious signs of toxicity, making 3-HPA a promising class for further development as antimalaria agents.

In this study, we investigated the antibacterial activity of 3-HPA. To this end we tested the previously published **7** (**TKK129**) and the additional 3-HPA derivatives **4** (**BLK278**), **5** (**BLK280**), and **6** (**TKK088**) (Fig. 1) against several clinically relevant Gram-positive and Gram-negative bacteria in minimal inhibitory concentration (MIC) assays. MICs were compared to those of mefloquine, lumefantrine, and halofantrine. The bacterial response to compound exposure was investigated using gel-based proteomics in the Gram-positive model organism *B. subtilis* 168. The proteomic profiles were interpreted based on the comparison of proteomic responses (CoPR) and a proteomic response library reflecting well over a hundred antibacterial agents [23]. Furthermore, the cytotoxicity and pharmacokinetics of compounds **4** (**BLK278**), **5** (**BLK280**), and **6** (**TKK088**) were assessed in this study.

**Fig. 1.**
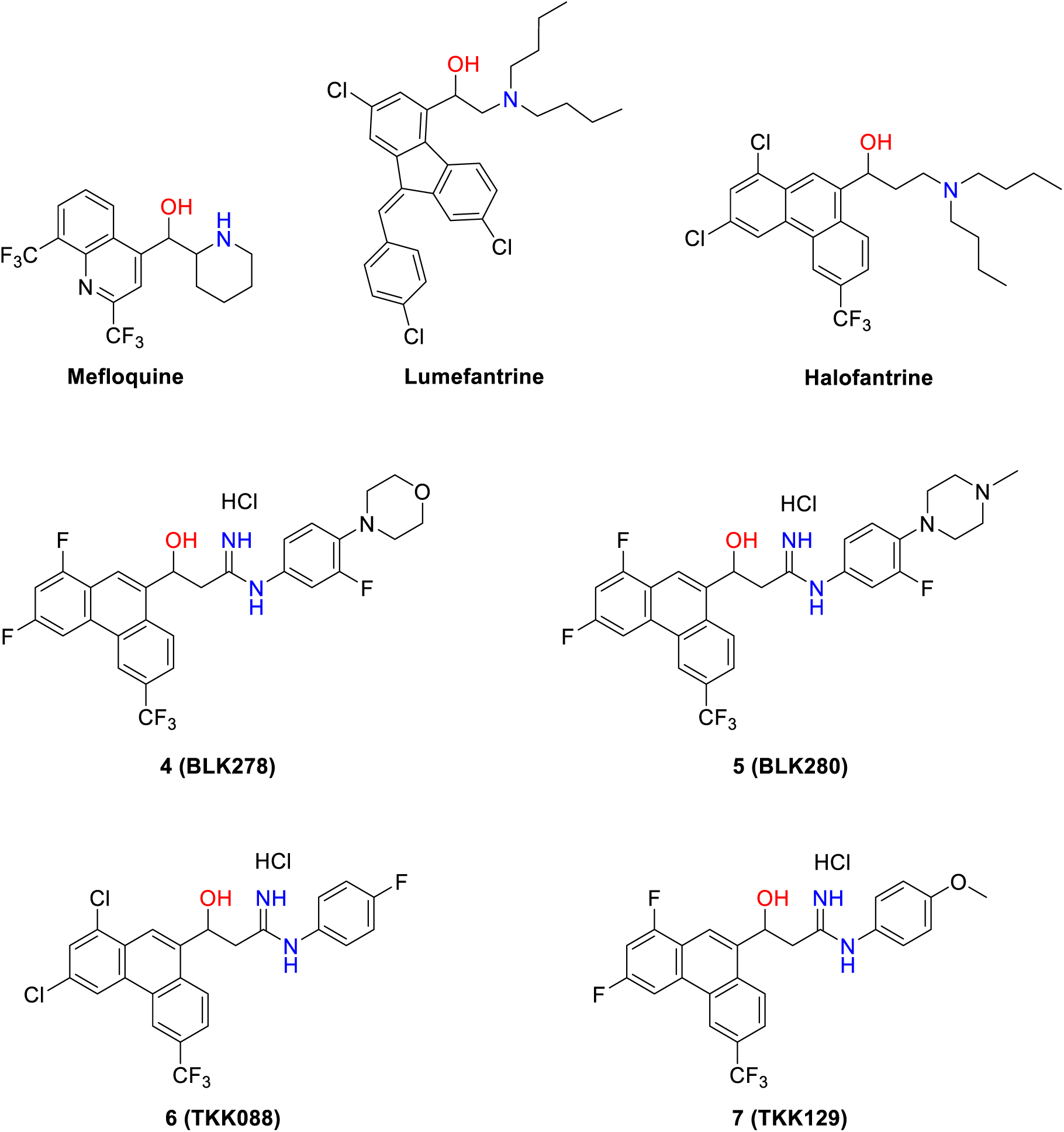
Structures of compounds investigated in this study. Mefloquine, lumefantrine and halofantrine are antimalarial arylamino alcohols included as reference compounds. The novel 3-hydroxypropanamidines **4** (**BLK278**), **5** (**BLK280**), and **6** (**TKK088**) were derived from the 3-HPA scaffold represented by the previously reported antimalarial lead **7** (**TKK129**) [21].

## Materials and methods

### Antibacterial activity

#### Minimal inhibitory concentration (MIC)

The MIC of all substances was determined (n = 2) in compliance with Clinical & Laboratory Standards Institute guidelines [24]. The procedure was performed as reported before, with the following modifications [25]. 1:2 dilution series were prepared utilizing the Fluent Automation Workstation (Tecan, Männedorf, Germany) ranging from 128–0.125 µg/ml. Mefloquine, halofantrine, lumefantrine, **4** (**BLK278**), **5** (**BLK280)**, **6** (**TKK088**) and **7** (**TKK129**) were tested against the following strains in Mueller-Hinton medium: *Bacillus subtilis* 168 (*trpC2*), S*taphylococcus aureus* DSM 20231 (ATCC 12600), *E. coli* BW25113, *E. coli* DSM 30083 (ATCC 11775), *E. coli* Δ*tolC*. Additional testing of derivatives against clinically relevant strains was performed in Mueller-Hinton medium, unless otherwise specified, and included: *Acinetobacter baumannii* DSM 30007 (ATCC 19606), *Enterobacter cloacae* DSM 30054 (ATCC 13047), *Klebsiella aerogenes* DSM 30053 (ATCC 13048), *K. pneumoniae* DSM 30104 (ATCC 13883), *Morganella morganii* DSM 30164 (ATCC 25830), *Proteus mirabilis* DSM 4479 (ATCC 29906), *Providencia rettgeri* DSM 4542 (ATCC 29944), *P. stuartii* DSM 4539 (ATCC 29914), *P. aeruginosa* DSM 50071 (ATCC 10145) (in cation adjusted Mueller-Hinton II medium), *Serratia marcescens* DSM 30121 (ATCC 13880), *Shigella sonnei* DSM 5570 (ATCC 29930), *S. aureus* ATCC43300 (MRSA)*, S. aureus* MU50.

### Cytotoxicity against human primary cells

Human umbilical vein endothelial cells (HUVEC) were purchased from PromoCell (Heidelberg, Germany) and cultured in Endothelial Cell Growth Medium (PromoCell). Human aortic smooth muscle cells (HAOSMC) were obtained from Innoprot (Derio, Spain) and cultured in Smooth Muscle Cell Medium-Plus (Innoprot).

Cells were seeded into 96-well plates at a density of 12,000 cells per well in 100 µl of the respective culture medium and incubated for 24 h at 37°C and 5% CO₂ to allow cell attachment. Subsequently, cells were exposed to the compounds for an additional 24 h under the same conditions. All compounds were dissolved in dimethyl sulfoxide (DMSO, Sigma-Aldrich), and the final DMSO concentration during treatment was 0.1% (v/v) in each well. Cells treated with 0.1% (v/v) DMSO alone served as vehicle controls. Final concentrations of mefloquine, halofantrine, **4** (**BLK278**), **5** (**BLK280**), **6** (**TKK088**), and **7** (**TKK129**) were 0.0625, 0.25, 1, 4, 16, 64, and 256 µg/ml, whereas lumefantrine was tested at 0.0625, 0.25, 1, 4, 16, and 64 µg/ml.

Thereafter, 10 µl of MTT stock solution (5 mg/ml; thiazolyl blue tetrazolium bromide; Gold Biotechnology, St. Louis, USA) was added to each well, followed by a further incubation period of 4 h. The resulting formazan crystals were dissolved in 100 µl DMSO per well, with gentle shaking for 15 min. Absorbance was subsequently measured at 570 nm using a Tecan Spark microplate reader (Tecan Group Ltd., Switzerland), with 630 nm used as the reference wavelength for background correction. Cell viability was calculated by normalizing the background-corrected absorbance of each compound-treated condition to that of the corresponding vehicle control, which was set to 100%.

IC₅₀ values were determined using GraphPad Prism version 10.6.1 for Mac by unweighted nonlinear least-squares regression of cell viability values against log₁₀-transformed concentrations, using a four-parameter logistic model with variable Hill slope. The upper plateau was fixed at 100%, while the lower plateau and Hill slope were fitted without constraints. IC₅₀ was defined as the concentration corresponding to the response halfway between the upper and lower plateaus.

### Determination of *in vitro* pharmacokinetics

Bioanalytical determination of **4** (**BLK278**), **5** (**BLK280**), and **6** (**TKK088**) was carried out on an Agilent 1200 Series HPLC system (Agilent, Waldbronn, Germany) coupled to a TSQ Quantum Ultra (Thermo Fisher Scientific, Waltham, Massachusetts, USA) mass spectrometer with electrospray ionization interface. The following transitions were monitored in positive ionization mode (MRM): 548.1 → 222.0 m/z for **4** (**BLK278**) (collision energy [CE]: 32 V, tube lens [TL]: 186 V), 561.1 → 234.0 m/z for **5** (**BLK280**) (CE: 31 V, TL:159 V), and 495.1 → 342.0 m/z for **6** (**TKK088**) (CE: 29 V, TL: 139 V). Chromatographic separation was achieved on a Waters XSelect CSH C18 3.0×150 mm, 3.5 µm (Waters, Eschborn, Germany). Using 0.1% formic acid in water (A) and methanol (B), the following gradient was applied at a flow rate of 400 µl/min: 0-1.5 min: 5% B, 1.5 – 8.0 min: 5 – 95% B, 8.0 – 12.0 min 95% B.

### Coefficient of distribution (logD)

The logD values were determined using the shake-flask method. An in-house protocol utilizing the Andrew+ pipetting robot (Waters Corporation, Eschborn) was applied [26]. Shortly, the distribution of 100 µM compound between 0.1 M potassium buffer pH 7.4 and octanol was investigated at three different buffer/octanol ratios and the volume ratio yielding the most similar distribution between both phases was used for calculation of the logD to ensure high accuracy. Carvedilol served as an internal control with published logD.

### Blood to plasma ratio

The ratio of the compounds between the red blood cell and plasma fraction was determined as described before [22] at two different concentrations: 50 nM and 1 µM (n=3). Briefly, the respective compound was spiked to fresh human whole blood, incubated for 30 min at 37 °C and plasma and red blood cells were separated by centrifugation (2000 x g, 10 min). The samples and corresponding references (compound added directly to plasma/red blood cells and incubated under the same conditions) were extracted with ice-cold acetonitrile. The extraction procedure was characterized by good recoveries (**4** 89.3%, **5** 91.2%, **6** 82.0%; n=3) and low matrix effects (**4** -10.7%, **5** -20.3%, **6** -18.1%; n=3). Obtained supernatant was dried under heated nitrogen stream (50°C), dissolved in acetonitrile/water (40:60 (v/v), and measured by LC-MS/MS. Carvedilol served as an assay control.

### Plasma protein binding

Protein binding was evaluated by equilibrium dialysis at a compound concentration of 50 nM and 1 µM [22]. Three independent replicates of compound-spiked plasma and pure 0.9% saline were placed in two chambers being separated by a regenerated dialysis membrane, and incubated for 24 hours at 37 °C. To enable more accurate determination of plasma protein binding exceeding 99%, plasma was diluted to 10% with 0.9% saline and incubated under the same conditions (“dilution method” [27]). Plasma was extracted by ice-cold acetonitrile in a ratio of 1:3 (v/v), evaporated and dissolved as described above. The saline samples from the acceptor chamber were supplemented with acetonitrile containing the internal standards to achieve a final acetonitrile:saline ratio of 2:3 (v/v), ensuring dissolution of the compounds. Analytes were quantified using matrix-matched calibration curves. Itraconazol served as assay control.

### Plasma stability

All compounds were tested as reported before [22] at two concentrations (50 nM and 1 µM) with samples taken after 0, 30, 60, 120, 240, 360 min, and 24 h in three independent replicates at 37 °C in human plasma. Furthermore, all three compounds were also spiked in 4% BSA to differentiate chemical and enzyme-related degradation. A compound was considered stable when the decrease from baseline did not exceed 15% (according to international bioanalytical guidelines) [28].

### Microsomal stability

The microsomal stability was investigated as described elsewhere [22] in human liver microsomes (HLM) pooled from 150 different adult donors of both sex (Corning, New York, USA). 1 µM **4** (**BLK278**), **5** (**BLK280**), and **6** (**TKK088**) were tested in biological triplicates at 37 °C. Reactions were stopped at 0, 15, 30, 45, and 60 min by the addition of acetonitrile. Negative controls (no NADPH cofactor) and blanks (no analyte), in addition to the positive control propranolol for cytochrome P450 enzyme activity, were included in each experiment. Half-life (t_1/2_) and intrinsic clearance (Cl_int_) were determined based on first-order kinetics.

### Microsomal protein binding

To account for non-specific binding in the HLM incubation, the binding of **4** (**BLK278**), **5** (**BLK280**), and **6** (**TKK088**) to HLMs was determined by equilibrium dialysis, following the procedure described above. However, plasma was replaced by a 0.5 mg/ml HLM suspension (without NADPH) in 0.1 M potassium phosphate buffer pH 7.4. The unbound intrinsic clearance, reflecting the enzymatic activity independent of non-specific binding, was calculated according to Equation 1.

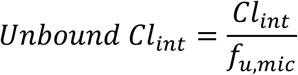

*Equation 1. Calculation of unbound intrinsic clearance. Clint: intrinsic clearance, fu,mic: unbound fraction microsomal incubation*

### Extrapolation of *in vivo* hepatic clearance

The *in vivo* hepatic clearance of compounds **4** (**BLK278**), **5** (**BLK280**), and **6** (**TKK088**) was predicted based on the well-stirred liver model encompassing the aforementioned blood-to-plasma ratio, unbound intrinsic clearance, and plasma protein binding (at 1 µM, according to Equation 2). The calculations were based on physiological parameters representative of a healthy adult male, including a hepatic blood flow of 1,350 ml/min, a microsomal protein content of 40 mg/g liver, and a liver weight of 1,500 g. Furthermore, the hepatic extraction ratio was determined according to Equation 3. Based on the calculated extraction ratios, compounds were classified as exhibiting low (<0.3), intermediate (0.3–0.7), or high (>0.7) hepatic extraction.

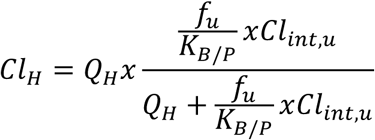

*Equation 2. Calculation of hepatic clearance. ClH: hepatic clearance, Clint,u: unbound intrinsic clearance, fu: fraction unbound in plasma, KB/P, blood-to-plasma ratio, QH: hepatic blood flow*

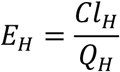

*Equation 3. Calculation of the hepatic extraction ratio (EH). ClH: hepatic clearance, QH: hepatic blood flow*

### Probing for bacteriolysis

A *B. subtilis* 168 culture was grown to an early logarithmic phase (OD_500_=0.35) in Belitzky Minimal Medium (BMM: 15 mM (NH_4_)_2_SO_4_, 8 mM MgSO_4_ x 7 H_2_O, 27 mM KCl, 7 mM Na_3_Citrat x 2 H_2_O, 50 mM Tris, 0.6 mM KH_4_PO_4_, 0.78 mM L-tryptophan, 2 mM CaCl_2_, 1 µM FeSO_4_, 10 µM MnSO_4_, 4.5 mM glutamic acid, 11.1 mM glucose) at 37 °C [29]. The culture was subsequently split and the sub-cultures treated with mefloquine, **4** (**BLK278**), **5** (**BLK280**), **6** (**TKK088**), and **7** (**TKK129**) at 5x the physiologically effective concentration. The PEC is defined as the concentration at which the growth is inhibited by 50-80%, in comparison to an untreated control. As positive control for cell lysis nisin was used at concentrations of 0.25 and 1.25 µg/ml. OD_500_ was measured every 30 min until an untreated control reached the stationary phase. Growth curves were recorded in three independent biological replicates.

### Gel-based proteomic analysis

Sample generation, isoelectric focusing, SDS PAGE, and protein identification were performed as described before [30] with minor modifications. Briefly, *B. subtilis* 168 cultures were grown in BMM to early logarithmic phase and then either left untreated as a control or treated with: 12.5 µg/ml mefloquine, 3.5 µg/ml **4** (**BLK278**), 6.5 µg/ml **5** (**BLK280**), 1.6 µg/ml **6** (**TKK088**), or 3.25 µg/ml **7** (**TKK129**), representing the physiologically effective concentrations. After 10 min of treatment with the respective compound all sub-cultures, including the control, were radioactively pulse-labelled with L-[^35^S]-methionine for 5 min. Translation - and pulse labelling - were terminated by adding chloramphenicol and a surplus of L-methionine. The cells were harvested, lysed, and the cytosolic proteome isolated. After determining the protein concentration (Nanoquant, Roth, Karlsruhe, Germany), approximately 50 µg radiolabelled (analytical gels) or 300 µg unlabelled (preparative gels) protein were used for isoelectric focusing. For the isoelectric focusing 24 cm immobilized pH gradient strips pH 4-7 (Immobiline DryStrip Gel, Cytiva, Uppsala, Sweden) were used. The separation of the proteome by molecular weight (second dimension) was performed with SDS-PAGE (40% acrylamide/bis solution, 29:1 (Bio Rad, Hercules, USA)). The resulting 2D gels were dried and autoradiographs documented (Typhoon Trio+, GE Healthcare, Uppsala, Sweden). The autoradiographs were analysed by creating overlays of images of treated samples with the respective controls, with the software Delta2D (4.8.2 Decodon, Greifswald, Germany). Based on the overlays, upregulation factors for specific spots were determined. Upregulation by a factor ≥ 2 defined marker spots (representing one or more proteins), which were cut out from preparative gels for protein identification by LC/MS.

Protein identification was performed as reported previously [31] with an ACQUITY UPLC I-Class / Vion IMS QToF MS^E^-system (Waters, Milford, Massachusetts, USA). Data was processed with the ProteinLynx Global Server (version 3.0.3, Waters) with the following parameters: lock mass (charge 1), 556. 2771 Da/e; Lock mass window, 0.25 Da; Low energy threshold, 50 counts; High energy threshold, 10 counts; Minimum number of fragments per peptide, 2; Minimum number of fragments per protein, 5; Minimum number of peptides per protein, 1; Maximum protein size, 25000; Digestive enzyme, trypsin; Skipped cleavages, 1; Fixed modifications, carbamidomethyl C; Variable modifications, oxidation M; False discovery rate, 4%. A *B. subtilis* 168 database (Uniprot ID: UP000001570) with added sequences for trypsin and keratin was used for protein identification. Data on spot identification were submitted to ProteomeXchange [32] via the PRIDE database [33]: project accession PXD080485 (Project Name: Proteomic Response of *Bacillus subtilis* towards mefloquine, **4** (**BLK278**), **5 (BLK280**), **6** (**TKK088**), and **7** (**TKK129**); Instruction for reviewers: please log in to PRIDE using the project accession PXD080485 and the token: VwOhJ5nTJ3oh).

Comparison of proteomic responses (CoPR analysis) was performed as described by Senges *et al*. [23]. In brief, to identify similar proteomic profiles in a reference compound library, the regulation factors of marker proteins identified by analysing autoradiographs of 2D gels with Delta 2D were logarithmised and used to perform cosine similarity calculations. Using the CoPR similarity matrix visualization was conducted by t-distributed stochastic neighbour embedding (t-SNE) (python v.3.8; perplexity = 7, learning rate = 500, 10000 iterations).

### L-[^35^S]-methionine incorporation assay

In parallel to the separation of cytosolic proteins, the protein samples were tested for L-[^35^S]- methionine incorporation to determine the impact of treatments on protein synthesis rates. To that end after determining the protein concentration with a Bradford assay (Nanoquant, Roth, Karlsruhe, Germany) 1 µl protein solution was spotted onto Whatman paper squares. After drying, samples were incubated in ice cold 20% trichloroacetic acid to precipitate proteins. The 20% trichloroacetic acid was discarded and two incubation steps in ice cold 10% trichloroacetic acid followed. After a final incubation with 96% ethanol at room temperature the paper squares were dried and placed into scintillation vials and radioactivity was measured with the Tri-Carb 2800TR scintillation counter (PerkinElmer). Measurements were performed in technical duplicates for all three biological replicates. Protein synthesis rates of the control were set 100% and used for normalization.

## Results

### Chemistry: Synthesis of amidines

A mini library of 3-hydroxypropanamidines (3-HPA) (Fig. 1) – three novel analogues (**4**-**6**) and one previously published 3-HPA (**7**) [21]– was synthesized with good yields and evaluated for antibacterial activity. 3-Hydroxypropanenitriles **1, 2** were used as starting materials and synthesized according to previously published methods [21]. In route A, the amidine moiety was introduced by reacting 3-hydroxypropanenitriles **1, 2** with the appropriate primary aromatic amine in the presence of trimethylaluminum, giving the corresponding novel 3-HPA **4-6** [34–36]. In route B, a Pinner reaction was used to convert the 3-hydroxypropanenitrile **1** into its imidoester hydrochloride **3**. Subsequent reaction with 4-methoxyaniline afforded **7** via aminolysis [21]. The ^1^H NMR and ^13^C NMR spectra are provided in the Supplementary Information (Suppl. Fig. S1-S9).

### 3-Hydroxypropanamidines show good antibacterial activity against bacterial model strains

The commercially available antimalarial agents halofantrine and lumefantrine were tested for antibacterial activity and compared to mefloquine, which acted as a control with already reported activity [3]. To test activity in Gram-negative bacteria three different *E. coli* strains were chosen. *E. coli* BW25113 is one of the commonly used laboratory strains and parental strain of the KEIO collection [37]. *E. coli* Δ*tolC* is an isogenic mutant with higher sensitivity to antibiotic treatment. This sensitivity is caused by the deletion of *tolC*, an essential part of the efflux pump system, and a subsequent membrane instability [38]. Finally, the type strain *E. coli* DSM 30083 was chosen as a standard reference. Type strains serve as taxonomic references representing the baseline characteristics of each species. For Gram-positive bacteria the model organism *B. subtili*s168 and the pathogenic type strain *S. aureus* DSM 20231 were used.

Mefloquine showed moderate antibacterial activity (8-32 µg/ml) (Tab. 1) against all strains. This is comparable to former studies [9]. Halofantrine and lumefantrine did not inhibit the growth of any strain at the tested concentrations up to 128 µg/ml. In addition to the commercially available antimalarial agents and **7** (**TKK129**), the newly synthesized 3-HPA were tested. Only the derivative **5** (**BLK280**) showed good activity against all tested model strains. However, it was the only derivative with lower activity against *S. aureus* (16 µg/ml) than mefloquine. Its unique residue is the methyl-piperazine (R^2^) (Fig. 2). The highest activity against *S. aureus* was observed for **6** (**TKK088**) (1 µg/ml) (Tab. 1). Furthermore, we tested the activity of the 3-HPA against 13 different clinically relevant bacterial strains (Tab. 2). Tested strains, like *A. baumannii*, *K. pneumoniae*, or *S. aureus* (MRSA) are listed by the WHO as high or critical priority pathogens [39]. All 3-HPA had good activity against the Gram-positive *S. aureus* strains, while activity against Gram-negative strains was generally limited. The broadest activity spectrum against Gram-negative strains was that of **5** (**BLK280**). Of note, **5** (**BLK280**) was active not only against the efflux-competent laboratory strain *E. coli* BW25113 but also against clinically relevant reference strains such as *A. baumannii* DSM 30007. In contrast, **4** (**BLK278**), **6** (**TKK088**), and **7** (**TKK129**) were much more active against the *E. coli* Δ*tolC* strain than the parental strain *E. coli* BW25113, indicating a greater impact of efflux and/or envelope permeability on their antibacterial activity. The greatest difference in activity was observed between **4** (**BLK278**) and **5** (**BLK280**), suggesting that the methyl-piperazine group in **5** (**BLK280**) contributes to its activity against Gram-negative bacteria. In addition, the di-chloro-substituted analogue **6** (**TKK088**) showed the lowest MIC against the high-priority *S. aureus* MRSA and VISA strains (0.5 µg/ml) (Tab. 2). Taken together, both antibacterial potency and activity spectrum of the 3-HPA were strongly influenced by variation of the N-aryl substituent of the propanimidamide moiety, while modifications of the phenanthrene scaffold were largely limited to exchange of fluorine for chlorine. For the previously published compound **7** (**TKK129**), antibacterial potency was more than two orders of magnitude lower than its antiplasmodial potency [21].

**Fig. 2.**
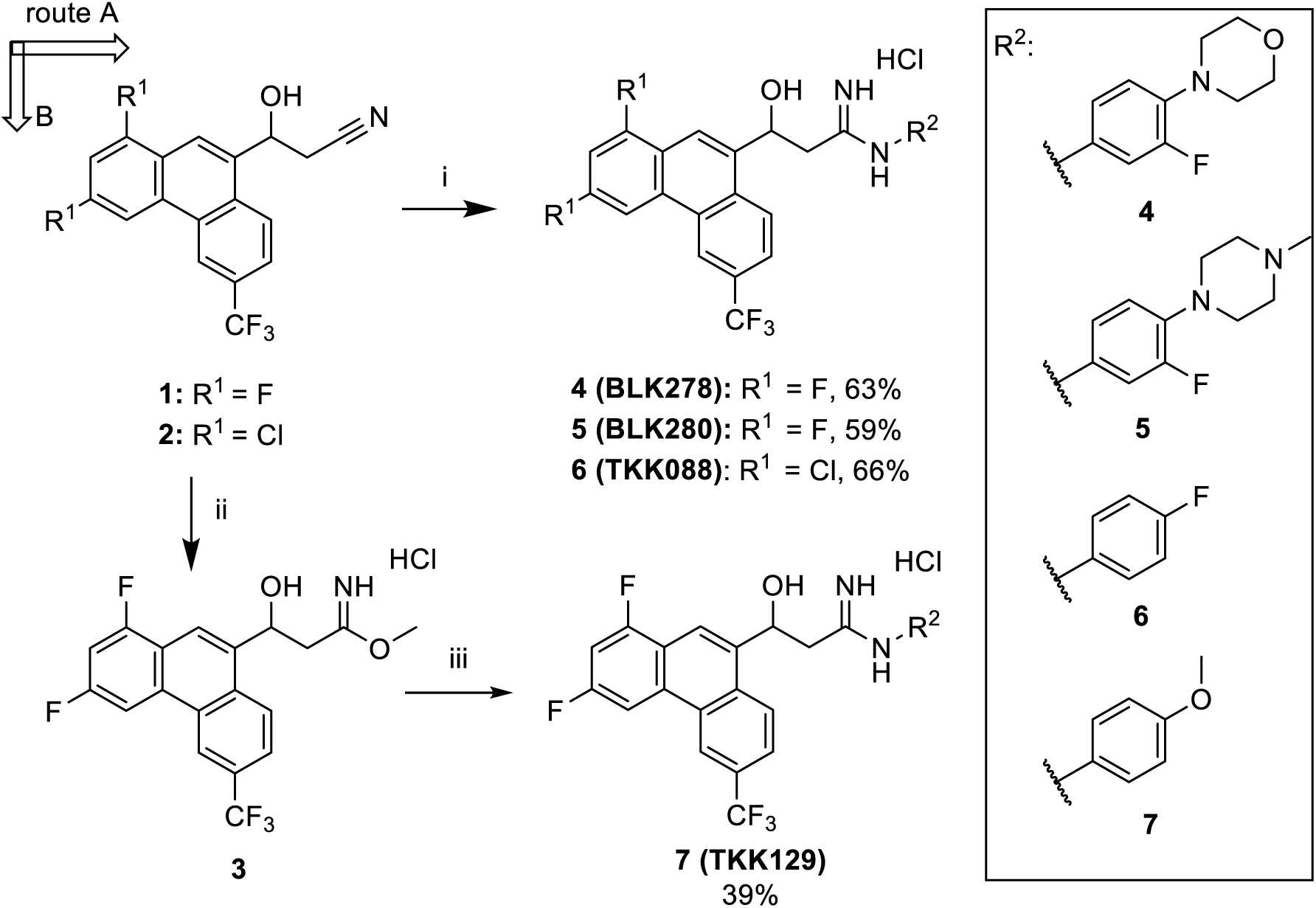
Synthesis of 3-HPA 4-6 (route A) and 7 (route B). (i) R^2^-NH_2_, Al(CH_3_)_3_, toluene/THF, 60°C, 24 h. (ii) MeOH, HCl in Et_2_O, CH_2_Cl_2_/THF, -10°C to RT. (iii) R^2^-NH_2_, DCM, 0°C to RT, 12 h [21].

**Tab. 1.**
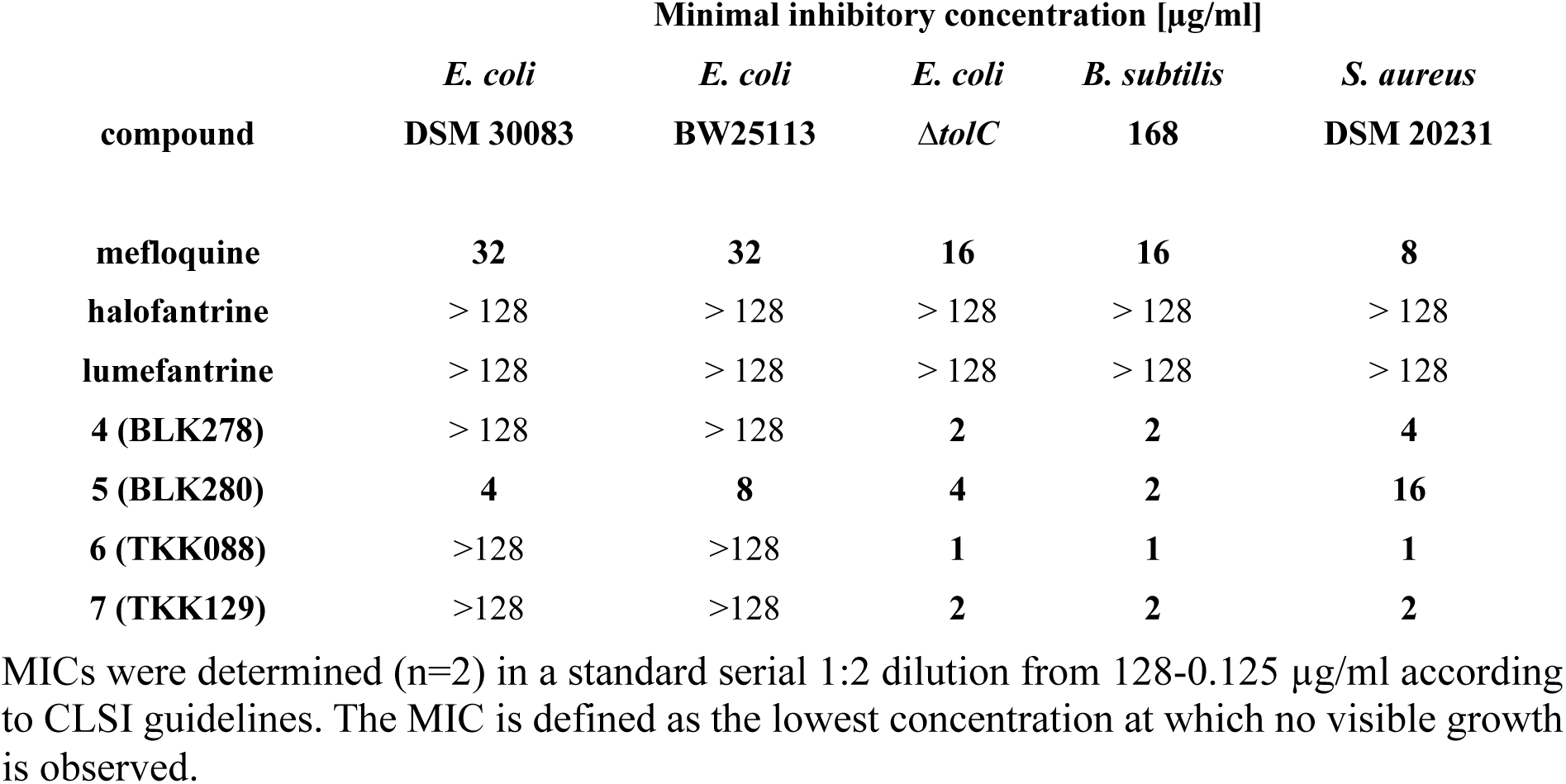
Minimal inhibitory concentrations (MIC).

| compound | Minimal inhibitory concentration [µg/ml] |  |  |  |  |
| --- | --- | --- | --- | --- | --- |
| | *E. coli*<br>DSM 30083 | *E. coli*<br>BW25113 | *E. coli*<br>$\Delta tolC$ | *B. subtilis*<br>168 | *S. aureus*<br>DSM 20231 |
| mefloquine | 32 | 32 | 16 | 16 | 8 |
| halofantrine | > 128 | > 128 | > 128 | > 128 | > 128 |
| lumefantrine | > 128 | > 128 | > 128 | > 128 | > 128 |
| **4 (BLK278)** | > 128 | > 128 | **2** | **2** | **4** |
| **5 (BLK280)** | **4** | **8** | **4** | **2** | **16** |
| **6 (TKK088)** | >128 | >128 | **1** | **1** | **1** |
| **7 (TKK129)** | >128 | >128 | **2** | **2** | **2** |

MICs were determined (n=2) in a standard serial 1:2 dilution from 128-0.125 µg/ml according to CLSI guidelines. The MIC is defined as the lowest concentration at which no visible growth is observed.

**Tab. 2.**
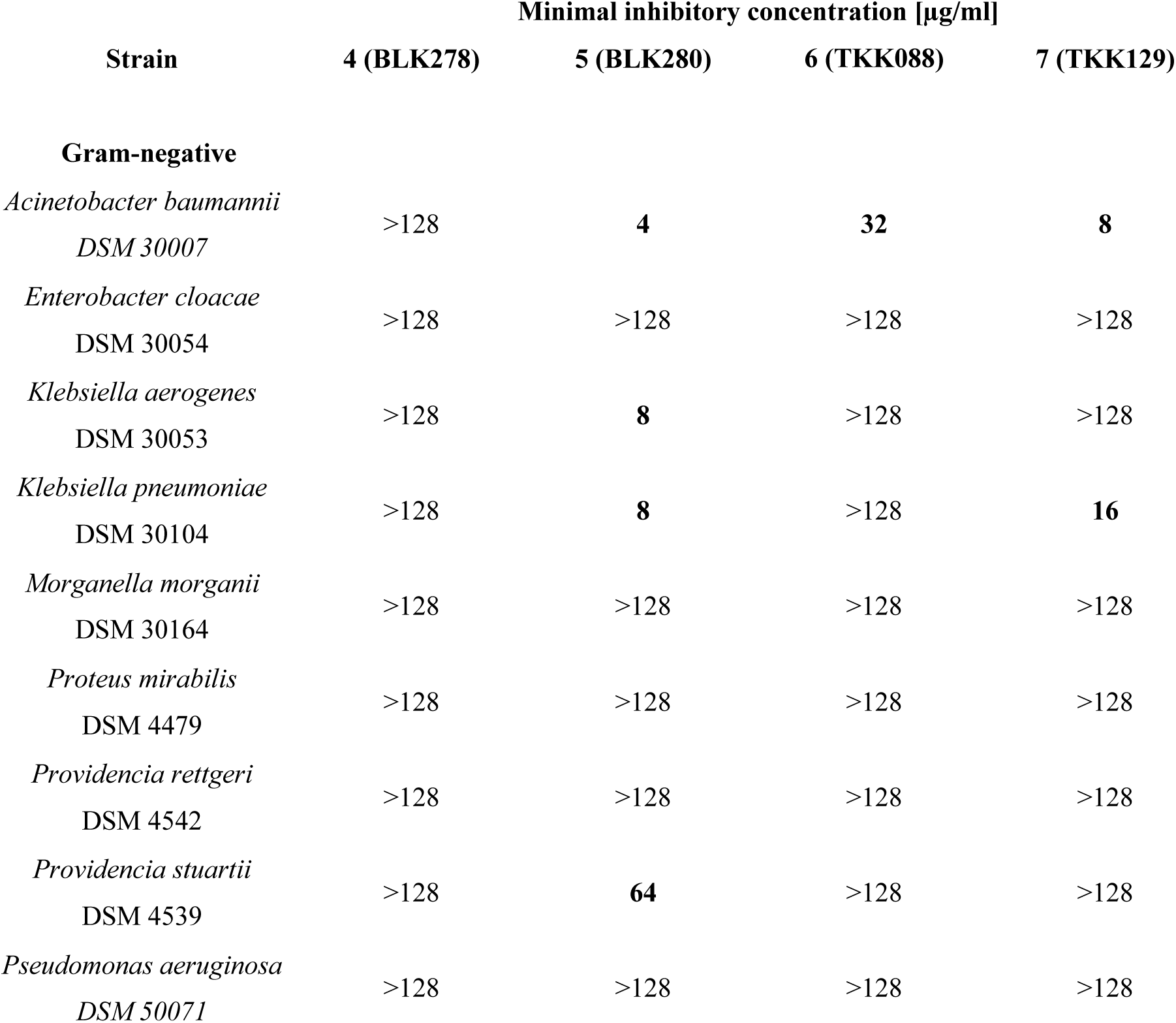
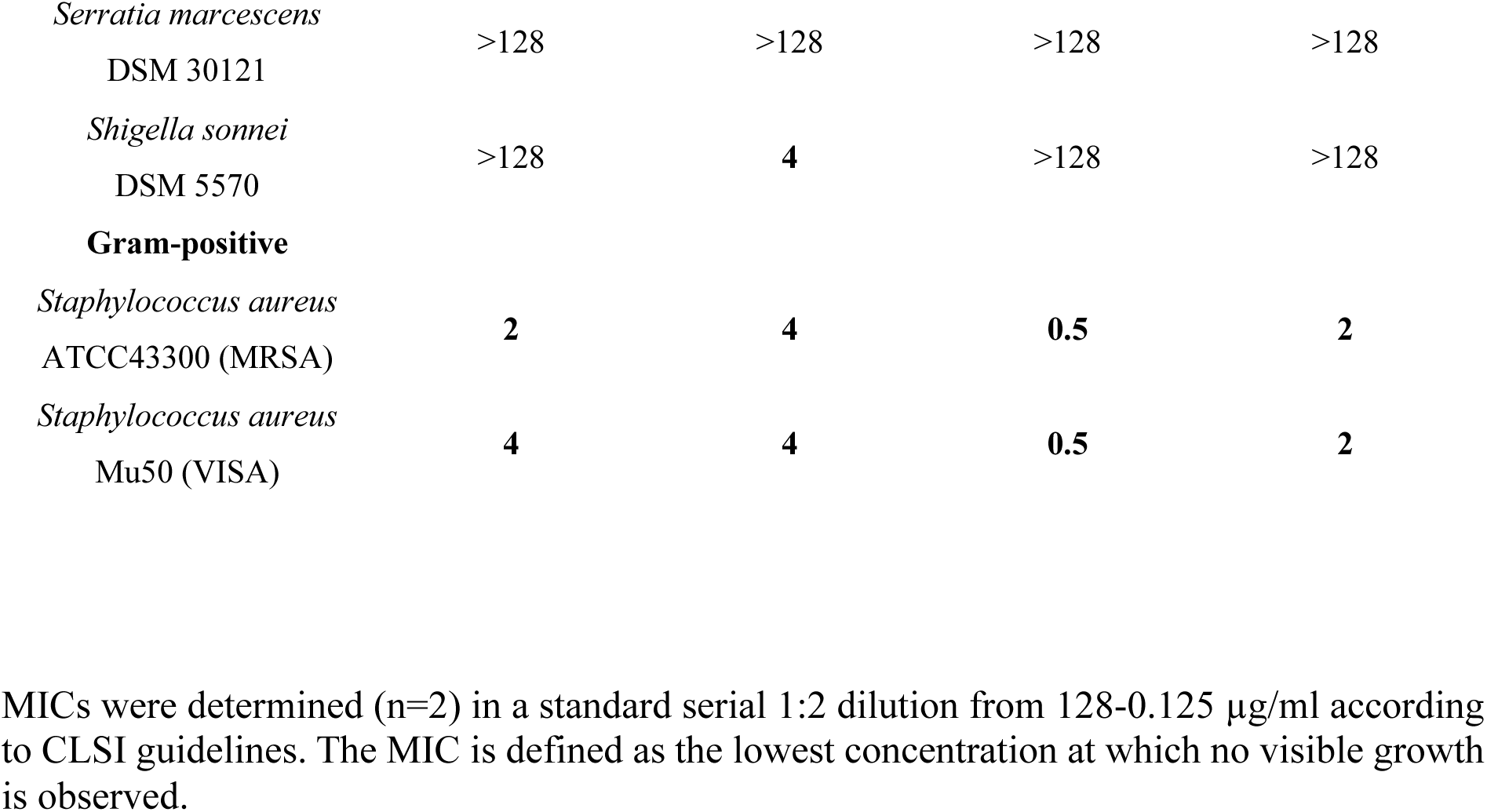
Minimal inhibitory concentrations (MIC) of four different 3-hydroxypropanamidines against clinically relevant Gram-positive and Gram-negative bacteria.

### *In vitro* cytotoxicity and selectivity of 3-hydroxypropanamidines

Cytotoxicity of the arylamino alcohols and 3-HPA was determined against human primary endothelial (HUVEC) and aortic smooth muscle (HAOSMC) cells by MTT assay (Tab. 3, Suppl. Fig. S10, Suppl. Fig. S11). All compounds reduced the viability of HUVEC and HAOSMC cells in the tested concentration range. Among the arylamino alcohols, halofantrine showed the lowest cytotoxicity in HAOSMC (IC_50_ values of 27.1 µg/ml), whereas lumefantrine was least cytotoxic in HUVEC (14.1 µg/ml). However, neither of these two compounds exhibited antibacterial activity at concentrations up to 128 µg/ml. IC_50_ values of the 3-HPA were at or up to an order of magnitude below those of mefloquine. Among the investigated 3-HPA, **4** (**BLK278**) and the previously reported **7** (**TKK129**) showed the most favourable overall cytotoxicity profiles, with cell-type-dependent differences. For **4** (**BLK278**) IC_50_ values were 1.27 µg/ml (1.71 µM) in HUVEC cells and 9.49 µg/ml (15.41 µM) in HAOSMC cells.

**Tab. 3.**
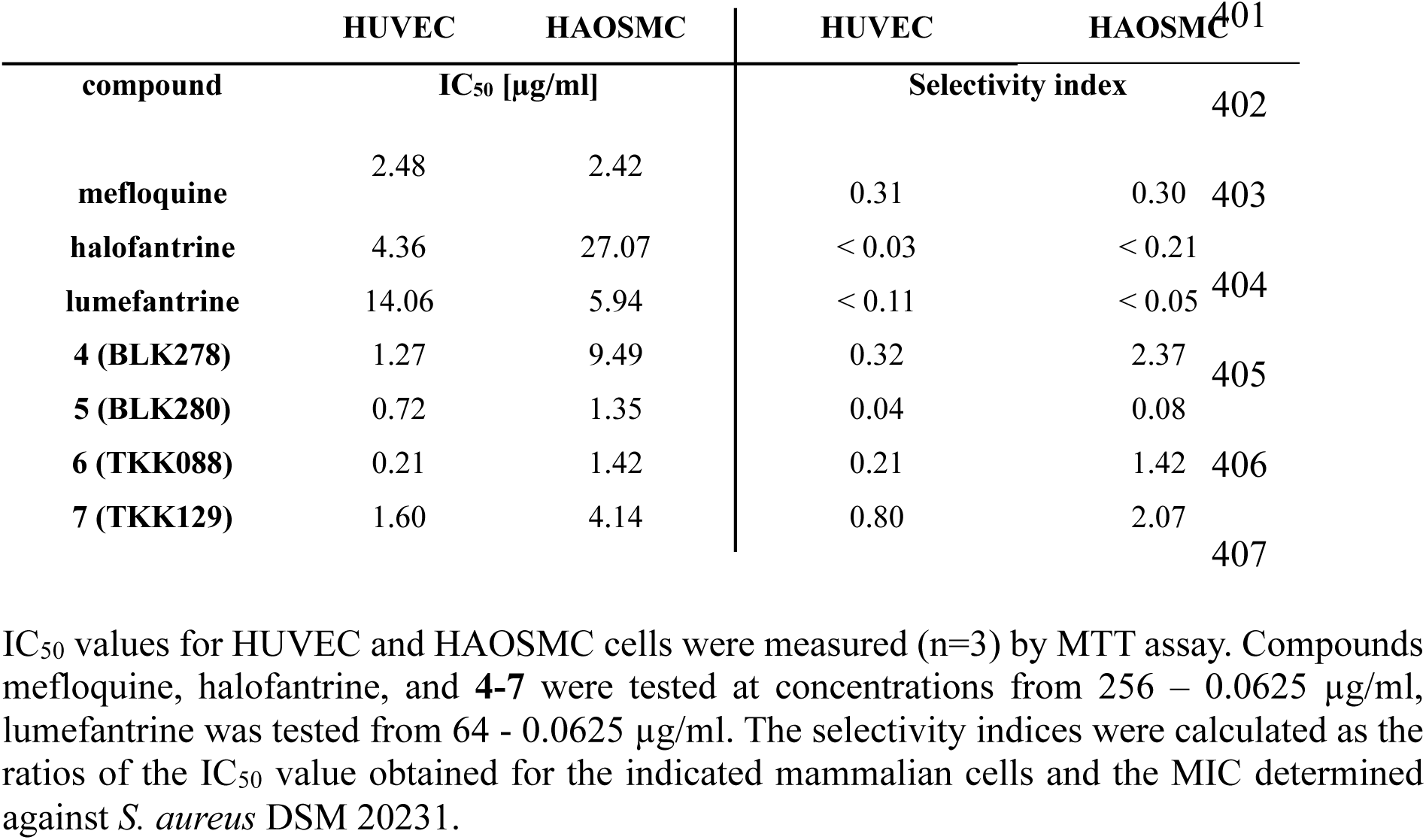
Cytotoxicity (IC_50_) and selectivity index (SI).

Taken together, the selectivity indices of the investigated arylamino alcohols and 3-HPA are considerably lower for antibacterial activity than for their established antiplasmodial activity, primarily reflecting the substantially higher concentrations required to inhibit bacterial growth. Importantly, the closely related lead compound **7** (**TKK129**) previously demonstrated excellent oral efficacy in a *P. berghei* infection model without obvious treatment-related toxicity [21]. Thus, while the *in-vivo* data support the general developability of the 3-HPA scaffold, improving antibacterial selectivity remains an important objective for further optimization.

### *In vitro* pharmacokinetics of the 3-hydroxypropanamidines 4 (BLK278), 5 (BLK280), and 6 (TKK088)

#### 3-Hydroxypropanamidines are characterized by high lipophilicity

The *in vitro* pharmacokinetics of **7** (**TKK129**) were reported elsewhere [21,22]. Here, we report the pharmacokinetics of **4** (**BLK278**), **5 (BLK280**), **6** (**TKK088**). The three newly characterized compounds exhibited high lipophilicity as determined from their measured distribution coefficient at pH 7.4 (logD_7.4_). LogD_7.4_ values were 4.73 ± 0.04 for **4** (**BLK 278**),

3.98 ± 0.11 for **5** (**BLK280**), and 5.20 ± 0.03 for **6** (**TKK088**), respectively (all n=3). These values indicate high lipophilicity typically associated with high passive membrane permeability. The control carvedilol confirmed assay validity with a value comparable to literature values (3.22 ± 0.04) [40].

#### 3-Hydroxypropanamidines exhibit high plasma protein binding

The binding to plasma proteins can limit the distribution and the metabolic availability of an antibiotic. At the same time, it can extend half-life and limit excretion [41]. All three 3-HPA exhibited high protein binding in plasma. Extent of protein binding correlated with compounds’ lipophilicity. **4** (**BLK278**) and **6** (**TKK088**) demonstrated plasma protein binding above 99%, while **5** (**BLK280**) displayed binding less than 99% (Tab. 4). All compounds demonstrated concentration-dependent plasma protein binding, with a lower unbound fraction observed at lower drug concentrations. For **7** (**TKK129**) >99% plasma protein binding was reported [21].

**Tab. 4.**
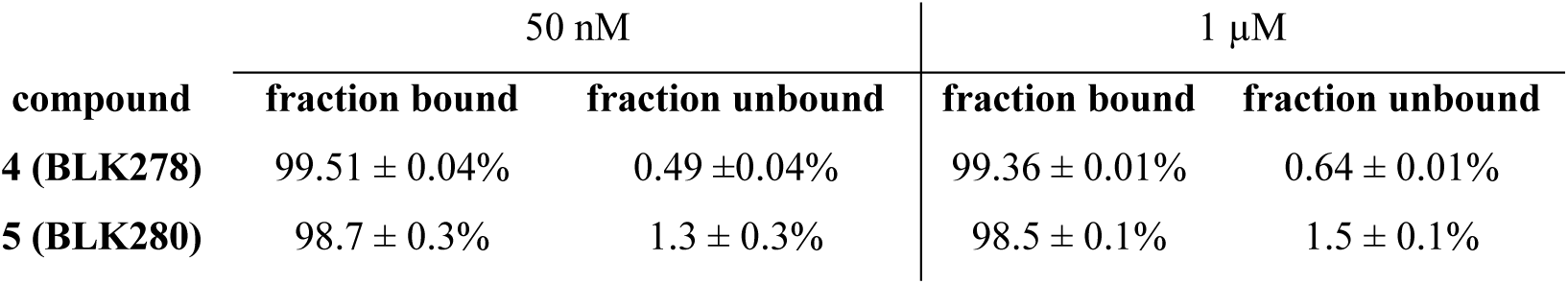
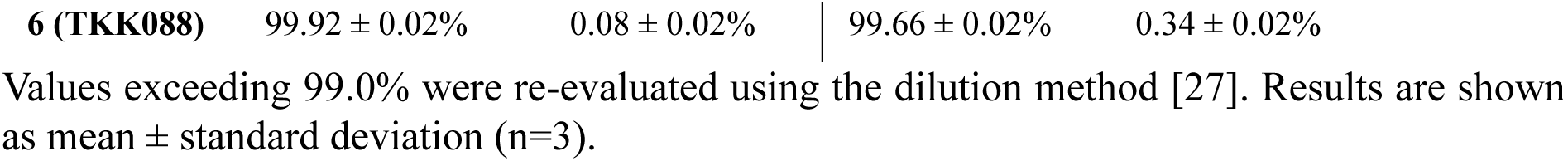
Plasma protein binding of the 3-hydroxypropanamidines.

#### 5 (BLK280) shows preferential distribution into whole blood

Blood-to-plasma (B/P) partitioning differed among the investigated compounds. Compound **5** (**BLK280**) exhibited the highest B/P ratio of >2, indicating preferential partitioning into blood cells, whereas compounds **4** (**BLK278**) and **6** (**TKK088**) distributed nearly equally in whole blood and plasma (Tab. 5). Similar distribution between blood and plasma was also reported for **7** (**TKK129**) with a ratio of 1.14 [21].

**Tab. 5.** Blood-to-plasma ratio of 3-hydroxypropanamidines.

| compound | 50 nM | 1 µM | Carvedilol (control) |
| --- | --- | --- | --- |
| **4 (BLK278)** | 0.931 ± 0.164 | 0.945 ± 0.031 | 0.649 ± 0.031 |
| **5 (BLK280)** | 2.934 ± 0.488 | 2.178 ± 0.181 | 0.633 ± 0.064 |
| **6 (TKK088)** | 0.951 ± 0.026 | 0.995 ± 0.048 | 0.620 ± 0.023 |

Blood-to-plasma ratios of **4 (BLK278)**, **5 (BLK280)**, and **6 (TKK088)**. Carvedilol served as a control substance with reported ratio. (n=3)

#### 3-Hydroxypropanamidines show excellent in vitro plasma stability

Plasma stability is another factor in drug distribution and availability. **4** (**BLK278**) and **5** (**BLK280**) remained stable in human plasma over 24 hours at both investigated concentrations (50 nM and 1 µM, Fig. 3A). **6** (**TKK088**) was likewise stable at 1 µM, whereas at a concentration of 50 nM a reduction of 18.8% was observed after 24 hours (Fig. 3A). Nevertheless, by taking typical *in vivo* half-life times into account, all three derivatives can be considered plasma stable compounds. Furthermore, **7** (**TKK129**) was also stable for more than 24 hours [22].

**Fig. 3.**
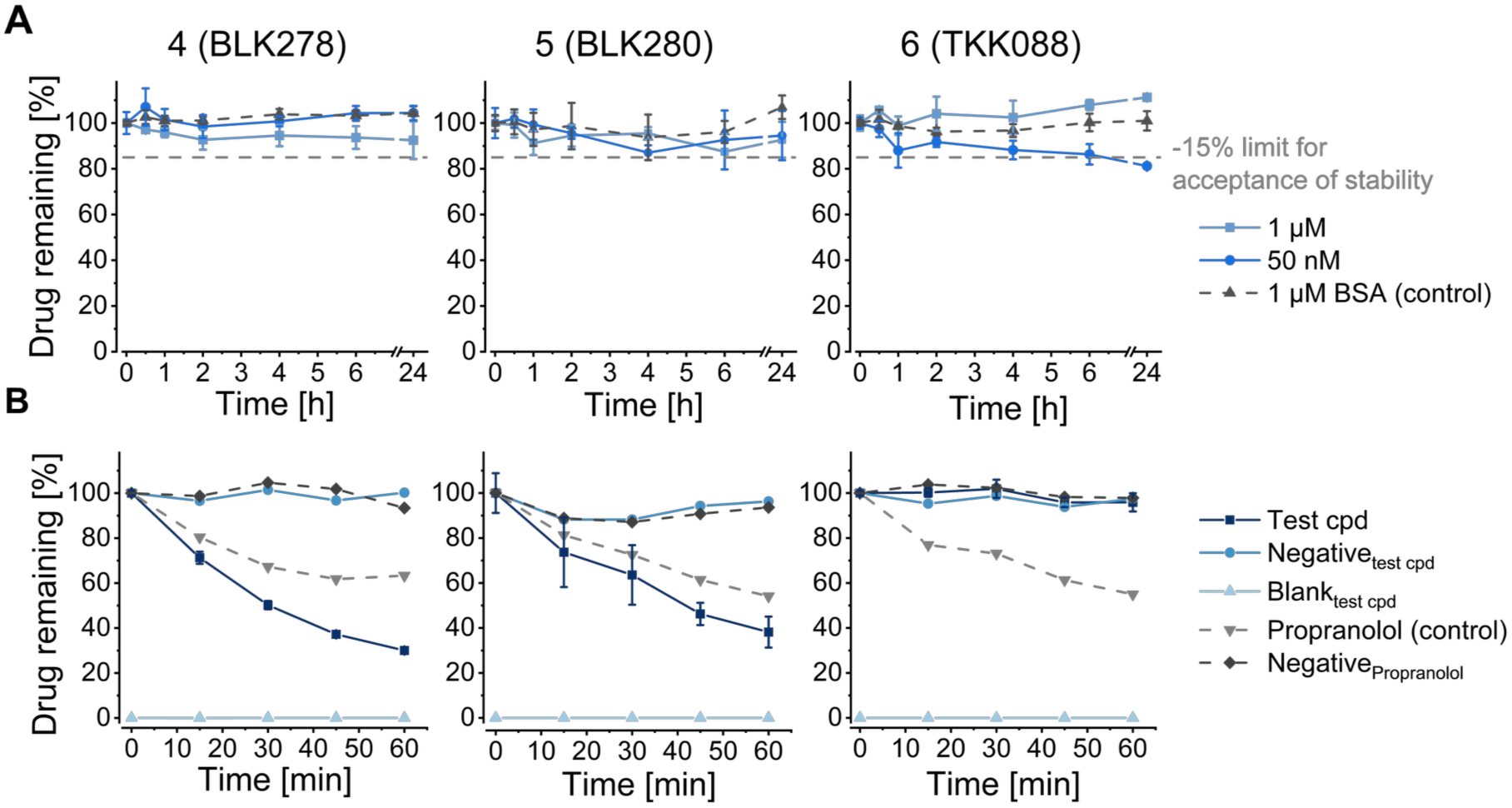
Stability of 3-hydroxypropanamidines in human plasma. **(A)** Plasma stability of **4** (**BLK278**), **5** (**BLK280**)**, 6** (**TKK088**). Incubation time was up to 24 h at 37 °C. A limit of - 15% was set as acceptance criterium for stability according to international bioanalytical guidelines (dashed grey line). 4% BSA (absence of plasma enzymes) was used as a control spiked with a concentration of 1 µM of the drug. **(B)** Microsomal stability of **4** (**BLK278**), **5** (**BLK280**), **6** (**TKK088**) in human liver microsomes. The CYP450 substrate propranolol was used as a control. The blank contains no drug, while the negative is free of cofactors necessary for CYP metabolism. Results represent mean ± standard deviation of n=3 independent replicates. BSA = bovine serum albumin, cpd: compound.

#### 3-Hydroxypropanamidines have intermediate to high microsomal stability

The microsomal stability of **4** (**BLK278**), **5** (**BLK280**), and **6** (**TKK088**) was evaluated in pooled human liver microsomes (Fig. 3B). Compounds **4** (**BLK278**) and **5** (**BLK280**) exhibited intermediate intrinsic clearance values of 40.8 ± 0.5 and 32.0 ± 2.5 µl/min/mg microsomal protein. In contrast, compound **6 (TKK088)** showed minimal turnover, with less than 15% degradation after 60 min of incubation. In line with guideline recommendations regarding low turnover rates, this finding was defined as a low intrinsic clearance (<5.4 µl/min/mg microsomal protein). For comparison, the reported intrinsic clearance for **7 (TKK129)** was 9 µl/min/mg (low) [21].

The investigated 3-HPA exhibited favourable *in vitro* pharmacokinetic properties, sharing several characteristics with the antimalarial amino alcohols mefloquine, halofantrine, and lumefantrine, while also showing compound-specific differences (Tab. 6). All compounds demonstrated high plasma protein binding and good plasma stability. Predicted hepatic extraction ratios were low for most derivatives, with the exception of compound **4** (**BLK278**), which showed an intermediate extraction ratio. Blood-to-plasma partitioning indicated similar distribution between blood and plasma for compounds **4** (**BLK278**) and **6** (**TKK088**), comparable to mefloquine [42] and **7** (**TKK129**) [22], whereas compound **5** (**BLK280**) displayed preferential partitioning into the blood compartment (B/P ratio >2). In contrast, halofantrine and lumefantrine showed a preference for the plasma compartment [42, 43]. The newly developed 3-HPA exhibited moderate to high lipophilicity, occupying an intermediate physicochemical space between mefloquine (logD_7.4_ 2.7) and the highly lipophilic antimalarial compounds halofantrine and lumefantrine (logD_7.4_ 5.68 and 7.34, respectively).

**Tab. 6.**
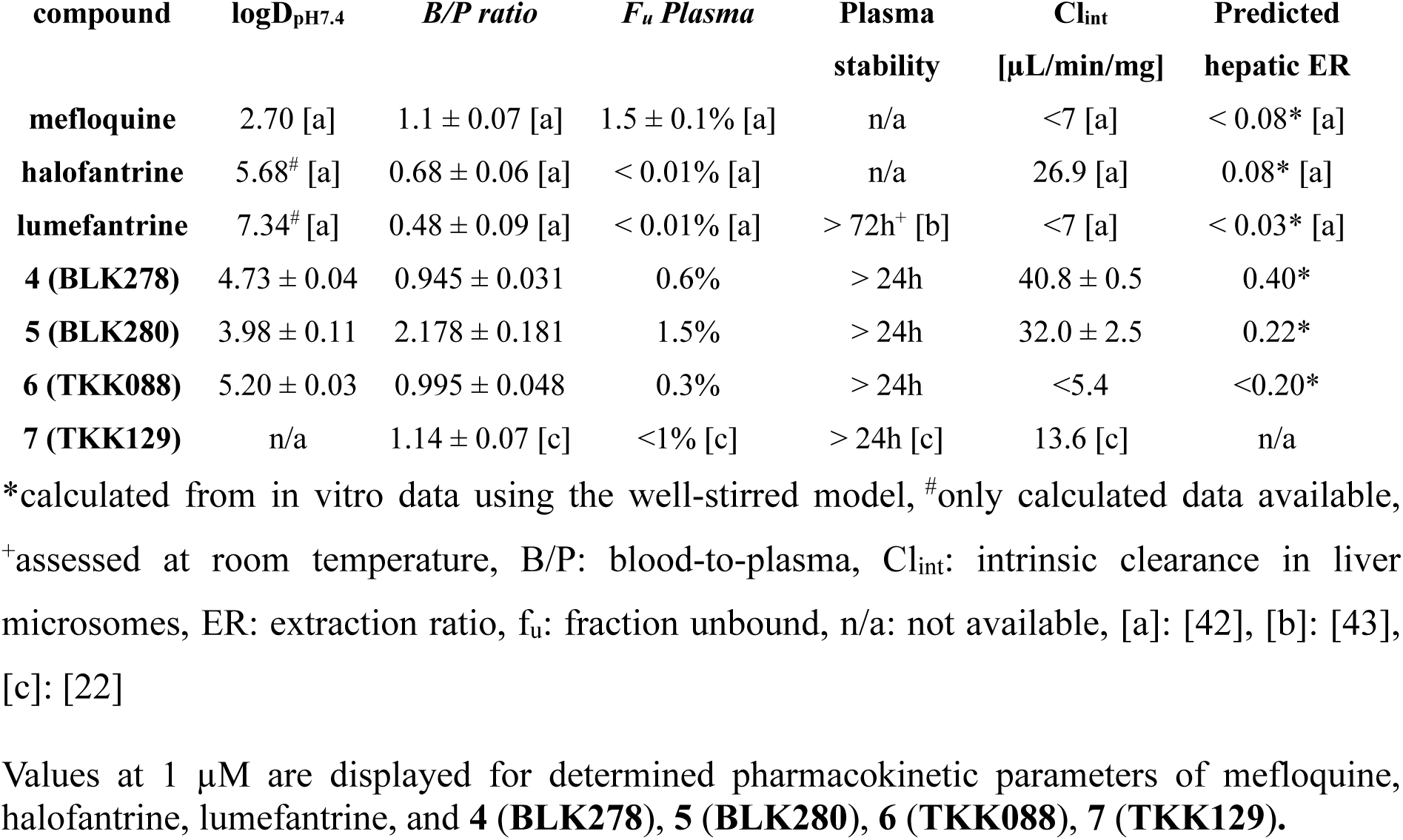
Comparison of *in vitro* pharmacokinetic characteristics.

### Impact of mefloquine and 3-hydroxypropanamidines on the model organism *B. subtilis*

The proteomic stress response of the Gram-positive model organism *B. subtilis* to mefloquine and the 3-HPA was analysed to gain an unbiased view of the cellular response to compound exposure. Growth experiments were performed to identify suitable concentrations for proteomic profiling that reduced growth rates by approximately 50% when applied in the early logarithmic phase - concentration we term physiologically effective concentration (PEC) (Fig. 4). All 3-HPA showed lower PEC than mefloquine (12.5 µg/ml). **6** (**TKK088**) (1.6 µg/ml) proved the most active. Interestingly, at five-fold the PEC, for all tested compounds a decrease in OD_500_ was observed. The data are consistent with bacteriolysis; however, the slower kinetics compared with nisin argue against rapid, nisin-like pore formation [44, 45].

**Fig. 4.**
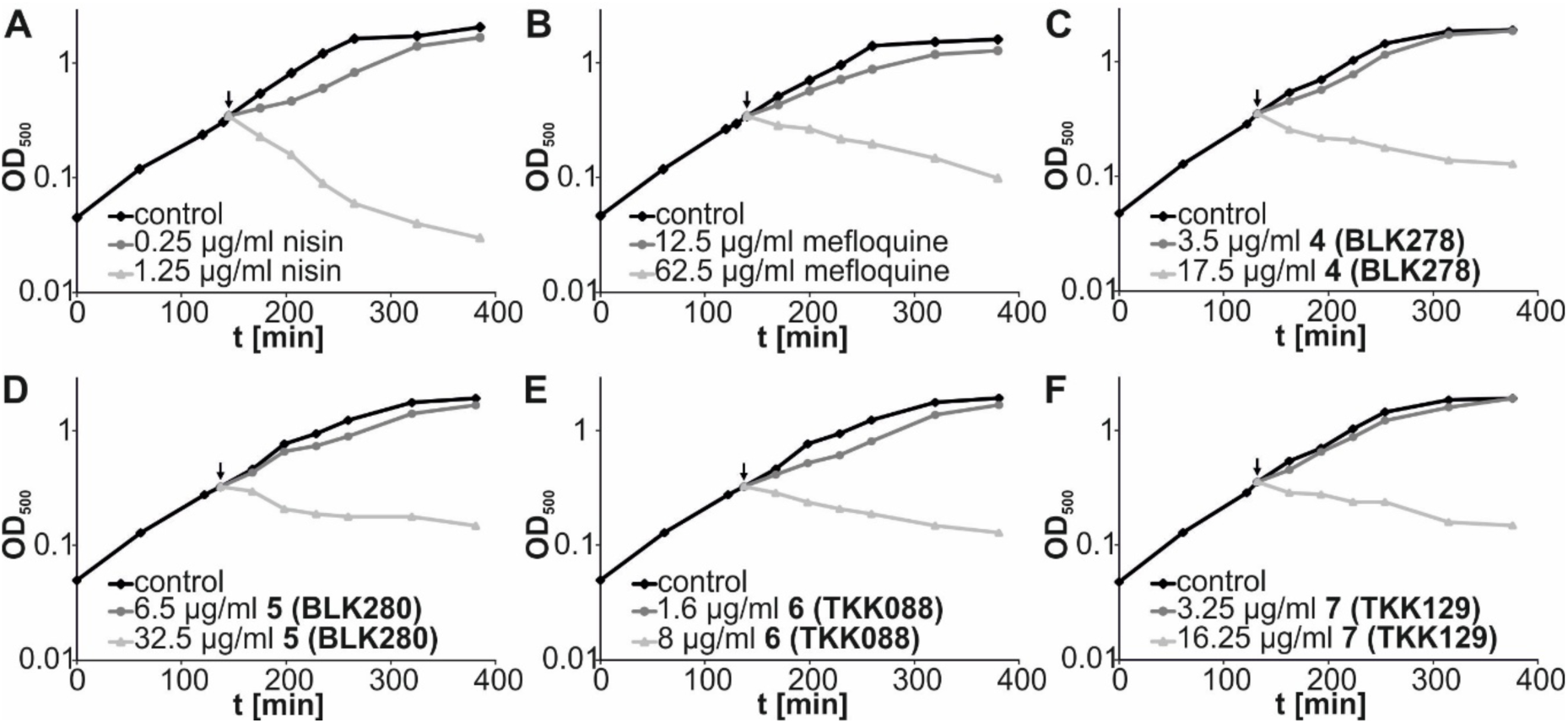
Effects on exponentially growing *B. subtilis*. Effects of different concentrations of nisin, mefloquine, **4** (**BLK278**), **5** (**BLK280**), **6** (**TKK088**), and **7** (**TKK129**), on *B. subtilis* growth in BMM medium. *B. subtilis* cultures were grown to an OD500 of 0.35 and either treated with the PEC or 5x PEC of the indicated agents or left untreated. **(A)** 0.25 or 1.25 µg/ml nisin, **(B)** 12.5 or 62.5 µg/ml mefloquine, **(C)** 3.5 or 17.5 µg/ml **4** (**BLK278**), **(D)** 6.5 or 32.5 µg/ml **5** (**BLK280**)**, (E)** 1.6 or 8 µg/ml **6** (**TKK088**), **(F)** 3.25 or 16.25 µg/ml **7** (**TKK129**). Arrows indicates the time of compound addition. Data shown are representative of three independent biological replicates. Note that **(C)** and **(D)** as well as **(E)** and **(F)** share the same controls.

To determine antibiotic-triggered changes in protein synthesis rates on a proteome-wide scale, L-[^35^S]-methionine was added to logarithmically growing *B. subtilis* cultures shortly after exposure to the antibiotic agents at the PEC. Relative protein synthesis rates were determined in relation to untreated controls to detect a potential impact on bacterial translation (Fig. 5). Neither mefloquine nor the tested 3-HPA reduced global protein synthesis rates at the tested concentrations 15 min after antibiotic addition.

**Fig. 5.**
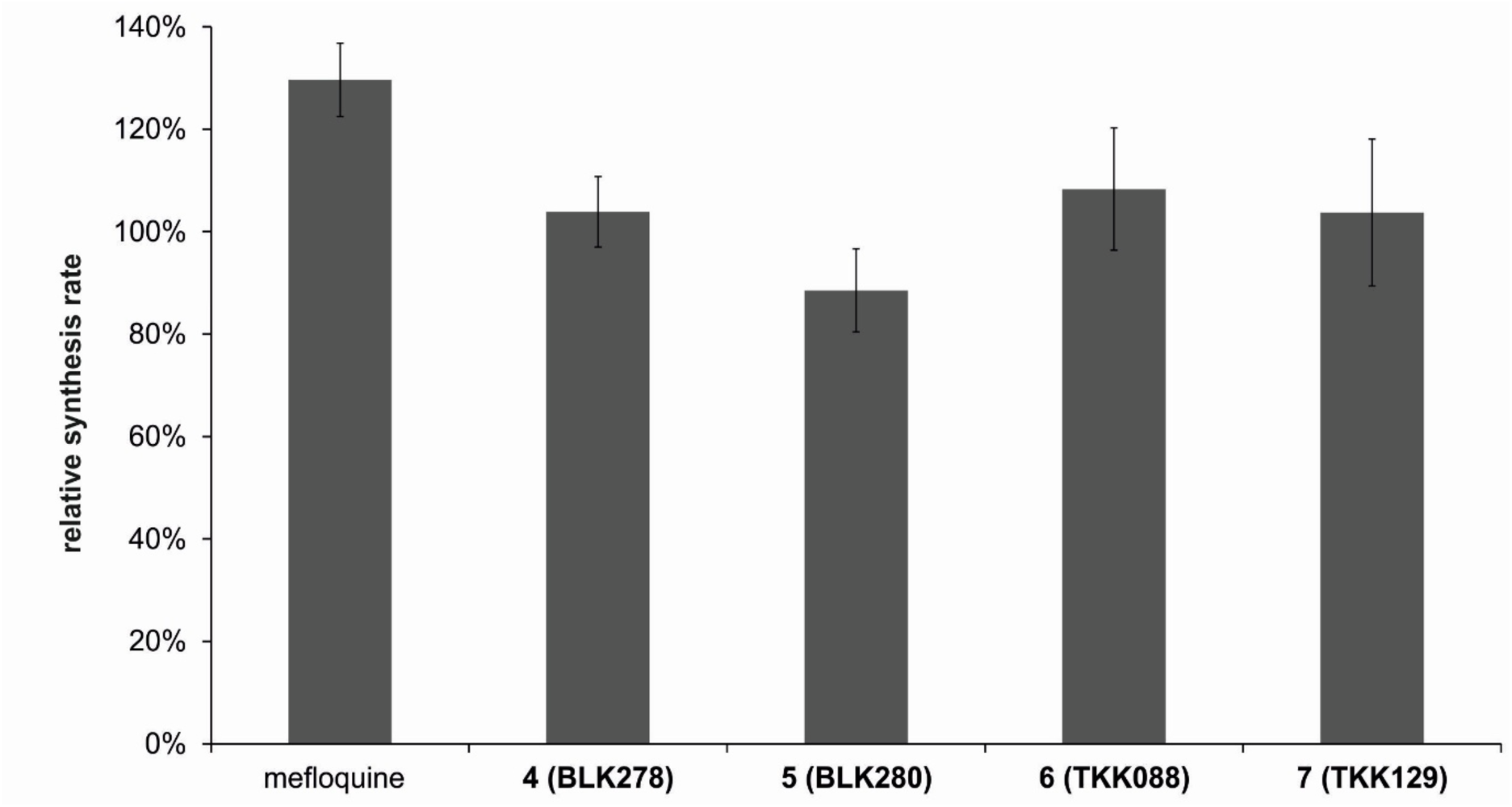
Effects on relative protein synthesis rate. Global relative synthesis rates of protein after treatment of *B. subtilis* with mefloquine and 3-HPA. Relative synthesis rates were determined by measuring the L-[^35^S]-methionine incorporation after pulse labelling (from min 10 to min 15 after compound addition) and normalizing to untreated controls. Radioactivity was measured in a scintillation counter. (n=3 with technical duplicates; means and standard deviations are given)

To assess protein synthesis rates of individual proteins, the proteomic responses of *B. subtilis* to mefloquine and **4** (**BLK278**), **5** (**BLK280**)**, 6** (**TKK088**), and **7** (**TKK129**) were analysed by 2D-gel-based proteomics and profiles compared to a published library of responses to antibiotics of different classes [23] (see Suppl. Fig. S12-S14 for representative growth curves). Overlays of autoradiographs of 2D gels representing cytosolic proteomes of untreated control and antibiotic-treated *B. subtilis* cultures, regulation factors and information on protein identification for each of the compounds can be found in the Supporting Information (Tab. S1-S10; Suppl. Fig. S15-S19). In the tSNE plot (Fig. 6) antibiotics cluster based on the similarity of their respective proteomic responses. The calculation of the similarity is based on proteins with upregulation factors ≥2 as described earlier [23, 46]. Mefloquine and the 3-HPA cluster together, indicating highly similar acute proteomic responses and suggesting similar effects on bacterial physiology. They cluster with substances known to interfere with membrane structure and membrane functions. Marker proteins indicative of impaired structural membrane integrity, membrane associated functions, and inhibition of cell wall biosynthesis are upregulated, alongside markers of proteotoxic and oxidative stress (Fig. 6). The observed changes are consistent with inhibition of the membrane-integral F₀-complex of the F₀F₁-ATPase, supporting F₀F₁-ATPase as a potential target of the 3-HPA, in line with previous findings for mefloquine in *S. pneumoniae* [9]. However, the proteomic responses also indicate pleiotropic effects, similar to those observed for the antimicrobial peptide MP196 [47], which clusters near mefloquine and the 3-HPA in the t-SNE plot (Fig. 6B). MP196 and gramicidin S have been shown to displace essential membrane-associated proteins such as MurG and cytochrome C [47].

**Fig. 6.**
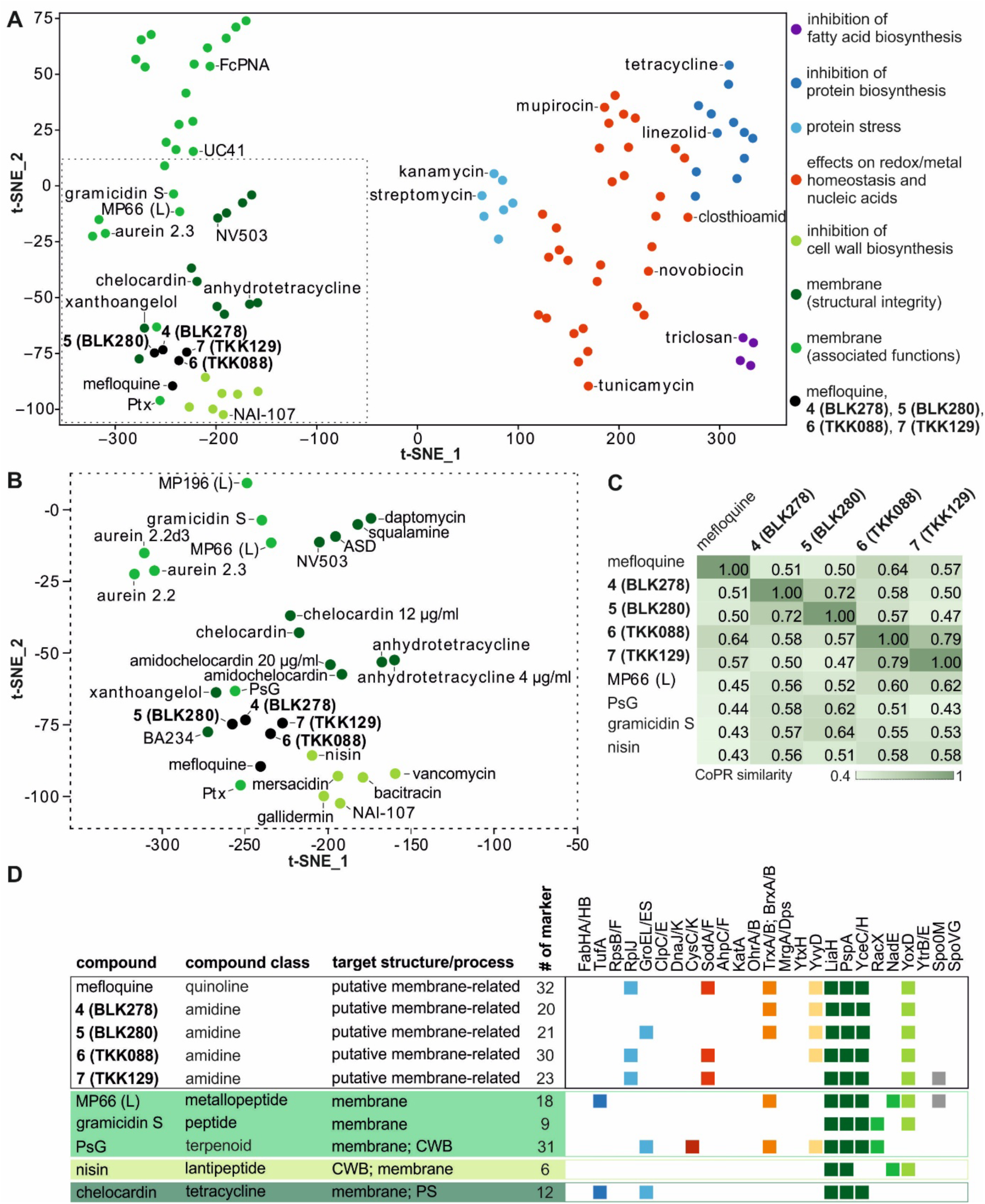
Proteomic response of *B. subtilis*. **(A)** t-SNE plot representing the similarity of *B. subtilis* proteomic responses to mefloquine and the 3-HPA in relation to more than 100 antibacterial agents with different modes of action. The used perplexity amounts to 7, the learning rate to 500 [23]. **(B)** Detail of the t-SNE plot focusing on mefloquine and 3-HPA. The colour code shown in **(A)** also applies to **(B)**. **(C)** Excerpt of the matrix of one-on-one similarity scores of the proteomic responses of *B. subtilis* to mefloquine, 3-HPA, and reference antibiotics [23]. **(D)** Similarity of proteomic responses to selected antibiotics that affect membrane targets. Squares indicate marker proteins informative of cellular structures or processes according to the following colour code: dark blue, protein synthesis; light blue, proteotoxic stress; dark red, sulphur metabolism; red detoxification of reactive oxygen species; orange, prevention of oxidative damage; yellow, general stress; dark green, membrane (structural integrity); medium green, membrane (associated functions); light green, inhibition of cell wall biosynthesis; grey, regulation of sporulation and cell division. PsG, pseudopterosin G; CWB, cell wall biosynthesis; PS, protein biosynthesis.

## Discussion

Mefloquine has been identified as an antibacterial agent, and investigations into the molecular target(s) and mode(s) of action were performed in different organisms and at differing depth. Published data converge in that the function of different membrane proteins in Gram-positive and Gram-negative bacteria were shown to be impaired [9–12]. Most notably, a mutation mapping approach revealed several mutations in the membrane-integral F_0_ subunit of the F_0_F_1_-ATPase in *S. pneumoniae* [9], which is instrumental in H^+^ channelling and thus ATP synthesis. In Gram-negative cells and mycobacteria, RND antibiotic efflux pumps or the mycolic acid transporter MmpL3 were implicated by cell-based assays [11–12] and *in silico* docking [10], respectively. Like F_0_F_1_-ATPase these proteins use the proton motive force, but it remains to be shown if they are direct targets of mefloquine action or affected by mefloquine-related changes of the membrane structure.

In this study, the broad activity spectrum of mefloquine was confirmed, with activities in the 8-32 µg/ml range (Tab. 1). Lumefantrine and halofantrine are somewhat larger molecules, which showed no antibacterial activity against the selected strains up to 128 µg/ml (Tab. 1). In earlier studies, similarly large heteroaromatic compounds showed higher activity against *S. pneumoniae* than mefloquine. In one of those compounds, OSU207, the quinoline group of mefloquine was exchanged for a phenanthrene group, with trifluoromethyl groups similar to the residues of halofantrine [9]. Here, we investigated the activity of three novel 3-HPA ((**4** (**BLK278**), **5** (**BLK280**), and **6** (**TKK088**)), together with the previously reported 3-HPA **7** (**TKK129**), which differ in their aromatic and amidine substituents. Within the 3-HPA series, antibacterial activity was influenced by both substitution of the phenanthrene core and variation of the amidine *N*-substituent. **6** (**TKK088**), bearing a 1,3-dichloro-substituted phenanthrene moiety, showed the highest activity against Gram-positive strains, whereas **5** (**BLK280**), bearing an N-methylpiperazine-containing substituent at the amidine nitrogen, displayed the broadest activity against Gram-negative strains. Notably, halogen substitution of the *N*-aryl substituent was not required for antibacterial activity, as the previously reported **7** (**TKK129**), bearing a 4-methoxyphenyl substituent, retained activity against Gram-positive and selected Gram-negative strains (Tab. 1, Tab. 2). It has previously been shown that amphiphilic compounds with aromatic groups often are good substrates for RND efflux pumps, especially those with multiple aromatic moieties [48]. Consistent with this, **4** (**BLK278**), **6** (**TKK088**), and **7** (**TKK129**) were markedly more active against the *E. coli tolC* deletion strain than against wild-type *E. coli* (Tab. 1). Because deletion of *tolC* also results in a general impairment of envelope integrity [49], these data cannot distinguish between effects on efflux and permeability In contrast, **5** (**BLK280**) showed much less dependence on TolC status and displayed the broadest spectrum of activity against Gram-negative pathogens (Tab. 2). Together, these findings indicate that efflux and/or envelope permeability are important determinants of the antibacterial activity of 3-HPA and that these properties can be modulated by variation of the amidine *N*-substituent.

For the previously reported compound **7** (**TKK129**), antibacterial potency was more than two orders of magnitude lower than its antiplasmodial potency [21]. In the present study, the compounds showed compound and cell-type-dependent cytotoxicity, with **4** (**BLK278**) and **7** (**TKK129**) displaying the most favourable profiles among the investigated 3-HPA (Table 3). These findings were consistent with the cytotoxicity previously reported for **7** (**TKK129**) in immortalized human cell lines [21]. *In vitro* pharmacokinetic profiling high plasma-protein binding, good plasma and microsomal stability of the 3-HPA. Regarding the blood-to-plasma ratio, **5** (**BLK280**) stands out with a higher affinity to red blood cells. The determined lipophilicity of 3-HPA may represent a more balanced profile, enhancing current solubility-related limitations of halofantrine and lumefantrine that require administration with food.

To gain an unbiased, global view of the impact of mefloquine and 3-HPA on bacterial physiology and to obtain potential clues to modes of action, a comparative proteome analysis was performed leveraging the proteomics response library for the Gram-positive model organism *B. subtilis* [23]. The proteomic responses were highly similar to one another so that they clustered together in the tSNE plot (Fig. 6). They also clustered with agents that target the cell envelope, such as pseudopterosin G, nisin, gramicidin S, or the positively charged small antimicrobial peptide MP196 and its derivatives, which - like gramicidin S - displace membrane-bound proteins [44, 46, 47, 50,51].

The most prominent marker proteins induced in mefloquine and 3-HPA-treated *B. subtilis* indicated an interference with the membrane structure and membrane-associated functions, but some indicators of proteotoxic stress and oxidative stress were also upregulated. For the parasite *P. falciparum* it has been shown that mefloquine, lumefantrine, and **7** (**TKK129**) inhibit heme detoxification, although the exact mechanism remains to be elucidated [52,21]. Given the marked difference between the antiplasmodial and antibacterial activities of 3-HPA, together with the absence of free heme in the growth medium of *B. subtilis* (BMM), inhibition of heme detoxification is unlikely to contribute substantially to the antibacterial activity observed under the present experimental conditions. For *Mycobacterium tuberculosis in silico* simulations predicted a membrane destabilizing effect of mefloquine most likely caused by a spacer effect of aromatic rings [53]. The dipeptide amide MC-207,110 caused an increased permeability of the outer membrane of *P. aeruginosa* PAM2035 [54]. Because the 3-HPA share key structural features with the arylamino alcohols, a similar membrane-destabilizing (“spacer”) effect may also contribute to their antibacterial activity.

In congruence with arylamino alcohols impacting the integrity and function of the bacterial cell envelope, *B. subtilis* cell lysis was observed at five-fold the physiologically effective concentration of mefloquine. The same effect was observed for 3-HPA. For *E. coli* a similar decrease in culture turbidity was reported earlier after mefloquine treatment [55], supporting a lytic effect on Gram-positive and Gram-negative bacteria. However, lysis of Gram-negative bacteria by the 3-HPA was not directly investigated here. For mefloquine, inhibition of translation was reported as a mode of action in *P. falciparum* [5]. Mefloquine was shown to bind directly to the GTPase-associated centre of the 80S ribosome, where the binding pocket is formed by ribosomal protein uL13 and 28S rRNA elements, including the eukaryote-specific expansion segment ES13 [5, 56]. This site is not structurally conserved in bacterial 70S ribosomes, which contain a 23S rRNA within the 50S subunit but lack eukaryotic rRNA expansion segments. Accordingly, no directly equivalent binding site is expected in *B. subtilis*, which is consistent with translation rates not being affected. However, upregulation of ClpP and GroEL indicate at least a mild proteotoxic stress, which could be related to impairment of membrane protein structure or function.

Taken together, the proteome analysis revealed a strong similarity between the responses to 3-HPA and mefloquine. Consistent with F₀F₁-ATPase being proposed as a potential target of mefloquine in *S. pneumoniae* [9], the proteomic response of *B. subtilis* indicated a pronounced impact on membrane structure and membrane-associated functions. Given the close resemblance of the proteomic profiles to those induced by gramicidin S and MP196, it is tempting to speculate that the 3-HPA have a similarly complex mode of action involving multiple membrane-associated processes. Multiple targets typically go along with low rates of spontaneous resistance.

## Conclusion

The investigated 3-hydroxypropanamidines exhibited antibacterial activity against clinically relevant Gram-positive pathogens, including MRSA and VISA, while selected analogues also showed activity against Gram-negative bacteria. All investigated 3-HPA were more active than mefloquine against the tested model strains *E. coli* Δ*tolC* and *B. subtilis*. Among the investigated analogues, **5** (**BLK280**) exhibited the broadest antibacterial spectrum, whereas **6** (**TKK088**) showed the highest activity against multidrug-resistant Gram-positive pathogens, namely *S. aureus* MRSA and VISA strains. Although the investigated 3-HPA exhibited limited selectivity toward primary human cells, their cytotoxicity was comparable to that of the clinically used arylamino alcohol antimalarials evaluated under identical experimental conditions. Comparative proteome analysis and cell lysis experiments suggest that interference with membrane-associated functions and cell envelope integrity contributes to their antibacterial activity. Although antibacterial selectivity remains limited, these findings identify the 3-HPA as a promising starting point for further structural optimization of antibacterial activity.

## Acknowledgements

The authors are grateful for excellent support by the technical staff at RUBION, Ruhr University Bochum. JEB gratefully acknowledges supported from the German Research Foundation (DFG) [RTG 2341 MiCon]. The use of the infrastructure of the “Center for System-based Antibiotic Research (CESAR)”, financed by the German Federal State of North Rhine-Westphalia and the European Union, European Regional Development Fund, Investing in your future (EFRE-0200598), is gratefully acknowledged.

## Supporting Information

Complete procedures for chemical synthesis and analysis and characterization data for all compounds are provided in the Supporting Information. The Supporting Information also contains information on identified proteins, supplementary pharmacological data, as well as supplementary data on the response of *B. subtilis* to compound treatment.

## Data availability

Additional data is shown in the supplementary material. Detailed data on protein identification is made available via the PRIDE repository: project accession PXD080485 (Project Name: Proteomic Response of *Bacillus subtilis* towards mefloquine, **4** (**BLK278**), **5** (**BLK280**), **6** (**TKK088**), and **7** (**TKK129**); Instruction for reviewers: please log in to PRIDE using the project accession PXD080485 and the token: VwOhJ5nTJ3oh).

