## Supporting Information for "3-Hydroxypropanamidines: from antiplasmodial leads to a novel antibacterial scaffold"

### Table of Contents

**Experimental Section (Chemistry)**

**Supplementary Tab. S1-S5** UniProt identifier, name and upregulation factor of marker proteins identified for *B. subtilis* 168 following treatment with mefloquine, **4** (**BLK278**), **5** (**BLK280**), **6** (**TKK088**), or **7** (**TKK129**)

**Supplementary Tab. S6-10** MS-based protein identification for mefloquine, **4** (**BLK278**), **5** (**BLK280**), **6** (**TKK088**), or **7** (**TKK129**)

**Supplementary Fig. S1-S9** $^{1}$H-NMR, $^{13}$C-NMR and HPLC spectra of novel 3-HPA

**Supplementary Fig. S10** Half-maximal inhibitory concentration ($IC_{50}$) of mefloquine, halofantrine, lumefantrine **4 (BLK278)**, **5 (BLK280)**, **6 (TKK088)**, and **7 (TKK129)** for HUVEC cells

**Supplementary Fig. S11** Half-maximal inhibitory concentration ($IC_{50}$) of mefloquine, halofantrine, lumefantrine **4** (**BLK278**), **5 (BLK280**), **6** (**TKK088**), and **7** (**TKK129**) for HAOSMC cells

**Supplementary Fig. S12** Inhibition of logarithmically growing *B. subtilis* 168 exposed to 12.5 µg/ml mefloquine

**Supplementary Fig. S13** Inhibition of logarithmically growing *B. subtilis* 168 exposed to 3.5 µg/ml **4** (**BLK278**) or 6.5 µg/ml **5** (**BLK280**)

**Supplementary Fig. S14** Inhibition of logarithmically growing *B. subtilis* 168 exposed to 1.6 µg/ml **6** (**TKK088**) or 3.5 µg/ml **7** (**TKK129**)

**Supplementary Fig. S15-S19** Proteomic response of *B. subtilis* 168 treated with 12.5 µg/ml mefloquine, 3.5 µg/ml **4 (BLK278**), 6.5 µg/ml **5** (**BLK280**), 1.6 µg/ml **6** (**TKK088**), 3.25 µg/ml **7** (**TKK129**)

**References**

#### Experimental section (Chemistry)

##### Chemicals and compounds

Mefloquine (Thermo Scientific), halofantrine (Merck) and lumefantrine (TCI) were purchased.

##### Chemical synthesis

##### General procedures for compound synthesis and analysis

($^1$H) and carbon ($^{13}$C) NMR spectra were recorded on a Bruker Avance 600, Bruker Avance NEO Evo − 600 (600.13 MHz for $^1$H and 125.76 MHz for $^{13}$C) and Bruker Avance 300 (300.13 MHz for 1H and 75.47 MHz for $^{13}$C) using DMSO-$d_6$ or MeOD as solvent. The coupling constants between two nuclei over n bonds ($^nJ$) are given in Hertz (Hz), while chemical shifts are presented in parts per million (ppm). All spectra were recorded at room temperature. Purity and chemical stability were determined by high-performance liquid chromatography (HPLC). A Knauer HPLC system in combination with a Vertex Plus column (150 × 4 mm with pre column, Eurospher II 100-5 C18) and a Knauer UV Detector K-2600 (Method 1) or UV-Detector Azura UVD 2.1L (Method 2) was used. The mobile phase 1 consisted of water with 0.1% trifluoroacetic acid in a linear gradient from 90–0%, the mobile phase 2 of acetonitrile with 0.1% of trifluoroacetic acid in a linear gradient from 10–100%. The total run time was 30 min, with a flow rate of 1 ml/min, followed by an isocratic elution with 100% acetonitrile over a period of 10 min. The purity of all compounds was determined with HPLC at 254 nm and scored > 95.0%. Melting points were measured automatically on a Cole-Parmer MP800 Series Stuart.

##### General procedures for synthesis of amidines

The starting materials (**1**, **2**, **3**) and **7 (TKK129)** were synthesized according to the literature as described by *Knaab et al.* [1]. Novel compounds **4 (BLK278)**, **5 (BLK280)**, and **6 (TKK088)** were synthesized in accordance with synthetic procedure **S1** outlined below.

##### General procedure for the synthesis of amidines using trimethylaluminum S1

Under an atmosphere of argon, the designated amine (2.0 equiv.) was dissolved in anhydrous toluene (5 ml/mmol) at room temperature. Trimethylaluminum (3.0 equiv., 2 M in toluene) was added and the solution stirred at 60°C for 30 min. Subsequently, a solution of 3-hydroxypropanenitrile **1** or **2** (1.0 equiv.) in anhydrous tetrahydrofuran (2.5 ml/mmol) and anhydrous toluene (2.5 ml/mmol) was added dropwise to the reaction mixture and stirred at 60°C for 16 h. Thereafter, the reaction mixture was poured onto an ice–water mixture (10 ml) and ethyl acetate (5 ml) were added. The phases were separated and the aqueous layer was extracted with ethyl acetate (3 × 20 ml). The combined organic phases were washed with saturated sodium chloride solution (3 × 20 ml) and dried over anhydrous $Na_2SO_4$. After filtration, the solvent was removed under reduced pressure, and the oily residue was dissolved in anhydrous diethyl ether. A 4 M solution of hydrogen chloride in dioxane was added dropwise to the ether phase while cooling to 0°C and the mixture was stirred for 5 min. The precipitate was subjected to repeated washings with acetonitrile, followed by drying and purification through flash chromatography [dichloromethane/methanol (10%)].

3-(1,3-Difluoro-6-(trifluoromethyl)phenanthren-9-yl)-*N*-(3-fluoro-4-morpholinophenyl)-3-hydroxypropanimidamide hydrochloride **4 (BLK278)**

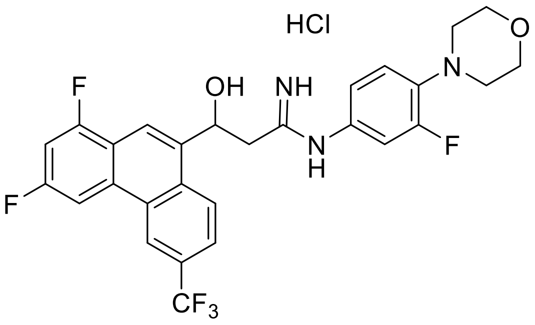


**4 (BLK278)** was synthesized according to the general procedure **S1**.

Brown solid, yield: 63%. mp: 217°C $^{1}$H NMR (600 MHz, DMSO-*d6*) δ 11.76 (s, 1H), 9.81 (s, 1H), 9.28 (s, 1H), 9.01 (d, *J* = 8.7 Hz, 1H), 8.84 (dd, *J* = 11.1, 2.3 Hz, 1H), 8.72 (s, 1H), 8.39 (s, 1H), 8.02 (dd, *J* = 8.7, 1.8 Hz, 1H), 7.72 (ddd, *J* = 10.9, 8.9, 2.3 Hz, 1H), 7.33 – 6.94 (m, 3H), 6.80 – 6.31 (m, 1H), 6.03 (dd, *J* = 10.2, 3.9 Hz, 1H), 3.77 (t, *J* = 4.6 Hz, 4H), 3.23 (dd, *J* = 13.7, 3.9 Hz, 1H), 3.06 (t, *J* = 4.7 Hz, 4H), 2.97 (dd, *J* = 13.7, 10.2 Hz, 1H); $^{13}$C NMR (151 MHz, DMSO) δ 164.75, 161.67 (d, J = 13.5 Hz), 160.04 (d, J = 13.6 Hz), 159.72 (d, J = 13.5 Hz), 158.06 (d, J = 13.8 Hz), 155.33, 153.70, 139.52, 1f39.47, 138.39, 131.81, 131.67, 128.91, 128.17, 128.11, 127.56, 127.35, 126.84, 125.27, 123.56, 123.47, 122.13, 122.00 (d, J = 2.8 Hz), 119.71 (d, J = 3.6 Hz), 117.52 (m), 113.82, 113.66, 105.63 – 105.32 (m), 103.58 – 103.04 (m), 67.18, 66.12, 50.32 (d, J = 3.2 Hz), 41.21; HPLC (Method 2): $t_R$ = 12.571 min, AUC = 99.0 %.

3-(1,3-Difluoro-6-(trifluoromethyl)phenanthren-9-yl)-*N*-(3-fluoro-4-(4-methylpiperazin-1-yl)phenyl)-3-hydroxypropanimidamide hydrochloride **5 (BLK280)**

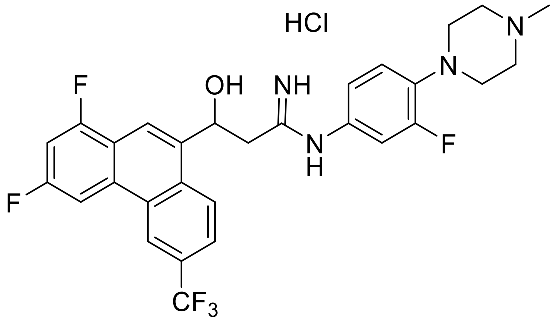


**5 (BLK280)** was synthesized according to the general procedure **S1**.

Brown solid, yield: 66%, mp: 247 °C; $^1$H NMR (600 MHz, DMSO-*d6*) δ 11.91 (s, 1H), 11.57 (s, 1H), 9.88 (s, 1H), 9.28 (s, 1H), 9.05 (d, *J* = 8.8 Hz, 1H), 8.92 – 8.65 (m, 2H), 8.40 (s, 1H), 8.01 (dd, *J* = 8.8, 1.8 Hz, 1H), 7.72 (ddd, *J* = 10.9, 8.9, 2.2 Hz, 1H), 7.38 – 7.01 (m, 3H), 6.58 (d, *J* = 4.7 Hz, 1H), 6.04 (d, *J* = 9.8 Hz, 1H), 3.50 (s, 4H), 3.35 – 3.08 (m, 5H), 2.98 (dd, *J* = 13.5, 10.2 Hz, 1H), 2.81 (s, 3H); $^{13}$C NMR (151 MHz, DMSO) δ 164.85, 160.10, 158.03, 154.48 (d, J = 246.7 Hz), 138.42, 138.17 (d, J = 9.0 Hz), 131.86, 131.68, 129.01, 128.94, 127.47 (q, J = 32.8 Hz), 126.89, 124.38 (d, J = 273.1 Hz), 123.67 – 123.52 (m), 122.14, 120.43, 117.75 – 117.42 (m), 113.90 (d, J = 23.2 Hz), 105.43, 103.27 (d, J = 26.3 Hz), 67.19, 52.21, 46.87, 41.97, 41.21; HPLC (Method 2): $t_R$ = 10.054 min, AUC = > 99.9%.

3-(1,3-Dichloro-6-(trifluoromethyl)phenanthren-9-yl)-*N*-(4-fluorophenyl)-3-hydroxypropanimidamide hydrochloride **6 (TKK088)**

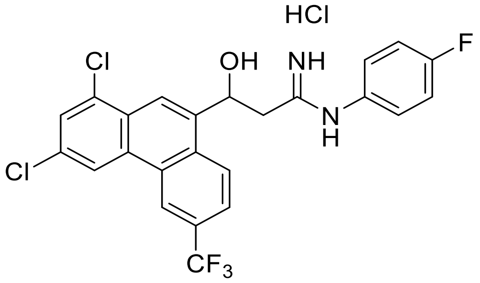


**6 (TKK088)** was synthesized according to the general procedure **S1**.

Off-white solid, yield: 66%, mp: 190 °C; $^1$H NMR (600 MHz, DMSO) δ 11.55 (d, *J* = 10.8 Hz, 1H), 9.76 (s, 1H), 9.41 (s, 1H), 9.27 (d, *J* = 2.0 Hz, 1H), 8.93 (dd, *J* = 8.4, 4.0 Hz, 1H), 8.70 (s, 1H), 8.61 (s, 1H), 8.13 – 8.02 (m, 2H), 7.45 – 7.36 (m, 4H), 6.60 (s, 1H), 6.01 (d, *J* = 9.9 Hz, 1H), 3.25 (dd, *J* = 13.8, 3.7 Hz, 1H), 2.91 (dd, *J* = 13.8, 10.2 Hz, 1H); $^{13}$C NMR (151 MHz, DMSO) δ 164.95, 162.19, 160.56, 140.37, 132.72, 132.25, 131.82, 131.24, 130.67 – 130.54 (m), 128.90, 128.18, 127.94 (d, *J* = 8.9 Hz), 127.83, 127.62, 126.99, 126.73, 125.26, 123.65,

123.45, 122.73, 122.20, 121.17, 116.95, 116.80, 67.16, 41.16; HPLC (Method 1): $t_R$ = 14.067 min, AUC = 97.1 %.

### Supplementary Tables

**Tab. S1** UniProt identifier [2], name, regulation factors with standard deviations (SDs) as well as proteins functions, regulator and functional categories of marker proteins identified for *B. subtilis* treated with mefloquine. Listed proteins and factors were used for CoPR analysis [3].

| UniProt identifier | Name | Regulation factor ± SD | Protein function | Regulator | Functional category |
|---|---|---|---|---|---|
| O32201 | LiaH | 48.69 ± 21.76 | subunit of the TatAY-TatCY protein secretion complex | LiaR | cell wall |
| P04990 | ThrC | 10.77 ± 8.88 | threonine synthase | Cody, ThrR | amino acids |
| O32078 | YuaE | 8.88 ± 1.31 | bacillithiol S-transferase | - | - |
| O05394 | MccB | 6.70 ± 1.62 | cystathionine lyase/homocysteine gamma-lyase | Spx, SigA, CymR | amino acids |
| O07609 | YhfK | 4.99 ± 1.70 | ene reductase | - | - |
| P94388 | Cah | 4.96 ± 1.72 | cephalosporin C deacetylase | - | cell wall |
| P54569 | YqkF | 4.88 ± 1.64 | NADPH-dependent 4-Hydroxy-2,3-trans-nonenal reductase | SigE | oxidative stress |
| P42923 | RplJ (RL10) | 4.54 ± 0.89 | ribosomal protein L10 | RplJ | translation |
| P54464 | YqeY | 4.41 ± 1.64 | unknown | - | - |
| P26907 | GsiB | 4.28 ± 0.78 | general stress protein | SigI, SigB | general stress |
| P80880 | TrxB | 4.13 ± 1.36 | thioredoxin reductase | Spx | oxidative stress |
| P14802 | YoxD | 3.90 ± 0.12 | similar to 3-oxoacyl- acyl-carrier protein reductase | - | cell envelope |
| P81100 | YceC | 3.77 ± 0.60 | similar to tellurium resistance protein | SigM, SigW, SigB, SigX | cell envelope |
| P54617 | PspA | 3.77 ± 1.11 | phage shock protein A homolog | SigW, AbrB | cell envelope |
| P21883 | PdhC | 3.64 ± 0.97 | pyruvate dehydrogenase | stringent response, SigA | energy |
| O34689 | MhqA | 3.19 ± 0.84 | hydroquinone-specific dioxygenase | SigA, MhqR | oxidative stress |
| P35137 | PpiB | 3.16 ± 0.53 | peptidyl-prolyl isomerase | SigA | protein stress |
| P28368 | YvyD | 3.10 ± 0.27 | ribosome hibernation promoting factor | SigB, PhoP, SigD, SigH | general stress, oxidative stress |
| P42976 | DapB | 3.06 ± 1.01 | dihydrodipicolinate reductase (NADPH) | Spx, SigA | cell wall |
| P54375 | SodA | 2.97 ± 0.97 | superoxide dismutase | SigB | oxidative stress |
| O05252 | NupN | 2.97 ± 0.52 | Lipoprotein, part of guanosine ABC transporter | SigA, CodY | nucleotide metabolism |
| P80864 | Tpx | 2.94 ± 0.57 | probable thiol peroxidase | Spx | oxidative stress |
| P54547 | G6pD | 2.73 ± 0.83 | glucose 6-phosphate dehydrogenase, pentose-phosphate pathway | - | energy |
| O32029 | YrrT | 2.67 ± 0.15 | putative AdoMet-dependent methyltransferase | Spx, SigA, CymR | - |
| P39645 | ChdC | 2.56 ± 0.71 | coproheme decarboxylase | - | - |
| O34305 | YtoQ | 2.52 ± 0.36 | unknown | - | - |
| O31669 | MtnD | 2.47 ± 0.31 | 1,2,-dihydroxy-3-keto-5-methylthiopentene dioxygenase | S-box, SigA | amino acids |
| O06724 | YisK | 2.44 ± 0.26 | oxaloacetate enol-keto tautomerase | Spo0Am SigH | cell envelope |
| Q45499 | AmpS | 2.35 ± 0.18 | aminopeptidase | - | proteolysis |
| P80244 | ClpP | 2.34 ± 0.52 | ATP-dependent Clp protease proteolytic subunit | CtsR, SigB, SigA | general stress. proteolysis |

| | | | | | |
|---|---|---|---|---|---|
| P80240 | GreA | 2.31 ± 0.50 | transcription elongation factor | - | transcription |
| Q06756 | IspF | 2.05 ± 0.06 | 2-C-methyl-D-erythrol-2,4-cyclodiphosphate synthase | SigM, SigB | general stress, lipid metabolism |

**Tab. S2** UniProt identifier [2], name, regulation factors with standard deviations (SDs) as well as proteins functions, regulator and functional categories of marker proteins identified for *B. subtilis* treated with **4** (**BLK278**). Listed proteins and factors were used for CoPR analysis [3].

| **UniProt identifier** | **name** | **Regulation factor ± SD** | **Protein function** | **Regulator** | **Functional category** |
|---|---|---|---|---|---|
| **O32201** | LiaH | 389.45 ± 175.22 | subunit of the TatAY-TatCY protein secretion complex | LiaR | cell wall |
| **P81100** | YceC | 6.18 ± 0.99 | similar to tellurium resistance protein | SigM, SigW, SigB, SigX | cell envelope |
| **P54524** | YqiG | 5.53 ± 1.08 | similar to NADH-dependent flavin oxidoreductase | | - |
| **P94400** | YciC | 5.37 ± 0.86 | ZTP activated GTPase A | SigA, Zur | metal ion homeostasis |
| **P54617** | PspA | 4.27 ± 1.09 | phage shock protein A homolog | SigW, AbrB | cell envelope |
| **O05394** | MccB | 4.15 ±1.26 | cystathionine lyase/ homocysteine gamma-lyase | Spx, SigA, CymR | amino acids |
| **O05239** | YugJ | 3.98 ± 0.59 | furan aldehyde reductase | Spx | - |
| **O34334** | YjoA | 3.89 ± 0.59 | bacillithiol S-transferase | - | - |
| **P46336** | IolS | 3.88 ± 0.35 | similar to dehydrogenase | SigA, IolR | carbon metabolism |
| **O07609** | YhfK | 3.6 ± 0.71 | ene reductase | - | - |
| **P14802** | YoxD | 3.59 ± 0.61 | similar to 3-oxoacyl- acyl-carrier protein reductase | - | cell envelope |
| **P32727** | NusA | 3.32 ± 0.98 | transcription termination factor | stringent response, SigA, SigB, NusA | transcriptiom |
| **P80880** | TrxB | 3.22 ± 0.58 | thioredoxin reductase | Spx | oxidative stress |
| **O34833** | YceH | 2.95 ± 0.42 | similar to toxic anion resistance protein | SigM, SigW, SigB, SigX | cell envelope |
| **P28368** | YvyD | 2.87 ± 0.45 | ribosome hibernation promoting factor | SigB, PhoP, SigD, SigH | general stress, oxidative stress |
| **P37573** | DisA | 2.85 ± 0.45 | DNA integrity scanning protein | SigM, Spx, SigA, SigB, CtsR, SigF | nucleotide metabolism |
| **P80244** | ClpP | 2.76 ± 0.47 | ATP-dependent Clp protease proteolytic subunit | CtsR, SigB, SigA | general stress, proteolysis |
| **O34305** | YtoQ | 2.45 ± 0.36 | unknown | - | - |
| **O07603** | YhfE | 2.25 ± 0.13 | similar to aminopeptidase | - | - |
| **P80874** | YhdN | 2.18 ± 0.21 | aldo/keto reductase | SigB, FapR | general stress, lipid metabolism |

**Tab. S3** UniProt identifier [2], name, regulation factors with standard deviations (SDs) as well as proteins functions, regulator and functional categories of marker proteins identified for *B. subtilis* treated with **5** (**BLK280**). Listed proteins and factors were used for CoPR analysis [3].

| **UniProt identifier** | **name** | **Regulation factor ± SD** | **Protein function** | **Regulator** | **Functional category** |
|---|---|---|---|---|---|
| **O32201** | LiaH | 205.33 ± 87.62 | subunit of the TatAY-TatCY protein secretion complex | LiaR | cell wall |
| **P54617** | PspA | 7.56 ± 1.88 | phage shock protein A homolog | SigW, AbrB | cell envelope |
| **P39605** | NfrA1 | 6.65 ± 0.91 | FMN-containing NADPH-linked nitro/flavin reductase | SigA, SigD, Spx, Spo0A | oxidative stress |
| **P54524** | YqiG | 5.99 ± 2.17 | similar to NADH-dependent flavin oxidoreductase | - | - |
| **P39645** | ChdC | 5.62 ± 1.46 | coproheme decarboxylase | - | - |
| **P28368** | YvyD | 4.65 ± 1.92 | ribosome hibernation promoting factor | SigB, PhoP, SigD, SigH | general stress, oxidative stress |
| **O05394** | MccB | 4.6 ± 0.80 | Cystathionine lyase/ homocysteine gamma-lyase | Spx, SigA, CymR | amino acids |
| **O05239** | YugJ | 4.37 ± 1.58 | furan aldehyde reductase | Spx | - |
| **P80880** | TrxB | 4.26 ± 0.85 | thioredoxin reductase | Spx | oxidative stress |
| **P46336** | IolS | 4.18 ± 1.00 | similar to dehydrogenase | SigA, IolR | carbon metabolism |
| **O34334** | YjoA | 3.99 ± 1.27 | bacillithiol S-transferase | - | - |
| **P80871** | YwrO | 3.94 ± 1.11 | similar to NAD(P)H oxidoreductase | - | - |
| **O30509** | GatB | 3.6 ± 0.46 | glutamyl-tRNA(Gln) amidotransferase (subunit B) | - | translation |
| **P81100** | YceC | 3.51 ± 0.28 | similar to tellurium resistance protein | SigB, SigM, SigW, SigX | cell envelope |
| **O34833** | YceH | 3.5 ± 0.79 | similar to toxic anion resistance protein | SigM, SigW, SigB, SigX | cell envelope |
| **P37967** | PnbA | 3.2 ± 0.40 | para-nitrobenzyl esterase | Abh, SinR, AbrB | lipid metabolism |
| **O07609** | YhfK | 3.18 ± 0.19 | ene reductase | - | - |
| **P14802** | YoxD | 3.07 ± 0.70 | similar to 3-oxoacyl- acyl-carrier protein reductase | - | cell envelope |
| **P80244** | ClpP | 2.72 ± 0.10 | ATP-dependent Clp protease proteolytic subunit | CtsR, SigB, SigA | general stress. proteolysis |
| **P80874** | YhdN | 2.66 ± 0.57 | aldo/keto reductase | SigB, FapR | general stress, lipid metabolism |
| **P28598** | GroEL | 2.52 ± 0.31 | Chaperonin and co-repressor for HrcA | SigA, HrcA | protein folding |

**Tab. S4** UniProt identifier [2], name, regulation factors with standard deviations (SDs) as well as proteins functions, regulator and functional categories of marker proteins identified for *B. subtilis* treated with **6** (**TKK088**). Listed proteins and factors were used for CoPR analysis [3].

| **UniProt identifier** | **Name** | **Regulation factor ± SD** | **Protein function** | **Regulator** | **Functional category** |
|---|---|---|---|---|---|
| O32201 | LiaH | 277.54 ± 145.67 | subunit of the TatAY-TatCY protein secretion complex | LiaR | cell wall |
| P54617 | PspA | 29.05 ± 27.00 | phage shock protein A homolog | SigW, AbrB | cell envelope |
| P31847 | YpuA | 14.16 ± 4.13 | unknown | SigM | - |
| O31711 | YknY | 8.42 ± 0.91 | ABC transporter | SigW, AbrB | cell envelope |
| P81100 | YceC | 5.87 ± 2.31 | similar to tellurium resistance protein | SigB, SigM, SigW, SigX | cell envelope |
| P02394 | RplJ (RL7) | 5.33 ± 0.28 | ribosomal protein bL12 | stringent response, RplJ, RplL | translation |
| P04990 | ThrC | 5.23 ± 1.22 | threonine synthase | Cody, ThrR | amino acids |
| O32078 | YuaE | 5.22 ± 1.30 | bacillithiol S-transferase | - | - |
| O05239 | YugJ | 5.05 ± 0.22 | furan aldehyde reductase | Spx | - |
| P39605 | NfrA1 | 5.03 ± 0.36 | FMN-containing NADPH-linked nitro/flavin reductase | SigA, SigD, Spx, Spo0A | oxidative stress |
| P14802 | YoxD | 4.46 ± 0.43 | similar to 3-oxoacyl- acyl-carrier protein reductase | - | cell envelope |
| P54569 | YqkF | 4.33 ± 1.19 | | | |
| P28368 | YvyD | 4.17 ± 1.39 | ribosome hibernation promoting factor | SigB, PhoP, SigD, SigH | general stress, oxidative stress |
| P39645 | ChdC | 3.98 ± 1.35 | coproheme decarboxylase | - | - |
| P42923 | RplJ (RL10) | 3.68 ± 0.42 | ribosomal protein L10 | RplJ | translation |
| O34969 | YfjR | 3.71 ± 1.67 | | | |
| P80244 | ClpP | 3.36 ± 0.71 | ATP-dependent Clp protease proteolytic subunit | CtsR, SigB, SigA | general stress, proteolysis |
| P38493 | Cmk | 3.22 ± 0.18 | cytidylate kinase | - | nucleotide metabolism |
| P23446 | FlgG | 3.20 ± 0.41 | flagellar hook protein | Spo0A, SigA, SinR, SwrA, SigD, CodY, DegU | - |
| O07607 | YhfI | 3.16 ± 0.47 | long-chain fatty-acid-CoA ligase | SigA, RoxS, FadR, CcpA | lipid metabolism |
| O31618 | ThiG | 3.09 ± 0.54 | thiamin thiazole synthase | Thi-box | - |
| P54464 | YqeY | 2.83 ± 0.75 | unknown | - | - |
| O32224 | AzoR2 | 2.79 ± 0.45 | similar to NAD(P)H dehydrogenase | SigA, SigG, MhqR | oxidative stress |
| P32396 | HemH | 2.77 ± 0.27 | coproporphyrin ferrochelatase | - | - |
| P37573 | DisA | 2.77 ± 0.19 | DNA integrity scanning protein | SigM, Spx, SigA, SigB, CtsR, SigF | nucleotide metabolism |
| P80864 | Tpx | 2.75 ± 0.18 | probable thiol peroxidase | Spx | oxidative stress |
| P54375 | SodA | 2.71 ± 0.32 | superoxide dismutase | SigB | oxidative stress |
| P80240 | GreA | 2.69 ± 0.37 | transcription elongation factor | - | transcription |
| P94424 | NfrA2 | 2.57 ± 0.22 | NADPH-FMN oxidoreductase | - | oxidative stress |
| O05394 | MccB | 2.43 ± 0.54 | cystathionine lyase/ homocysteine gamma-lyase | Spx, SigA, CymR | amino acids |

**Tab. S5** UniProt identifier [2], name, regulation factors with standard deviations (SDs) as well as proteins functions, regulator and functional categories of marker proteins identified for *B. subtilis* treated with **7** (**TKK129**). Listed proteins and factors were used for CoPR analysis [3].

| **UniProt identifier** | **Name** | **Regulation factor ± SD** | **Protein function** | **Regulator** | **Functional category** |
|---|---|---|---|---|---|
| O32201 | LiaH | 196.49 ± 167.60 | subunit of the TatAY-TatCY protein secretion complex | LiaR | cell wall |
| P71088 | Spo0M | 18.94 ± 5.54 | sporulation-control protein | SigW, SigH | sporulation |
| P54617 | PspA | 15.16 ± 19.20 | phage shock protein A homolog | SigW, AbrB | cell envelope |
| P31847 | YpuA | 14.91 ± 7.52 | unknown | SigM | - |
| P02394 | RplJ (RL7) | 5.84 ± 1.17 | ribosomal protein bL12 | stringent response, RplJ, RplL | translation |
| O05239 | YugJ | 5.68 ± 1.78 | furan aldehyde reductase | Spx | - |
| P39605 | NfrA1 | 4.77 ± 0.57 | FMN-containing NADPH-linked nitro/flavin reductase | SigA, SigD, Spx, Spo0A | oxidative stress |
| P04990 | ThrC | 4.70 ± 1.06 | threonine synthase | Cody, ThrR | amino acids |
| P54569 | YqkF | 4.64 ± 1.67 | NADPH-dependent 4-Hydroxy-2,3-trans-nonenal reductase | SigE | oxidative stress |
| P81100 | YceC | 3.77 ± 1.81 | similar to tellurium resistance protein | SigB, SigM, SigW, SigX | cell envelope |
| O32078 | YuaE | 3.57 ± 1.51 | bacillithiol S-transferase | - | - |
| P39645 | ChdC | 3.51 ± 1.03 | coproheme decarboxylase | - | - |
| P42923 | RplJ (RL10) | 3.47 ± 0.98 | ribosomal protein L10 | RplJ | translation |
| P49786 | AccB | 3.45 ± 0.42 | acetyl-CoA carboxylase (biotin carboxyl carrier subunit) | - | lipid metabolism |
| O05240 | YugK | 3.34 ± 0.78 | similar to NADH-dependent butanol dehydrogenase | - | - |
| P94424 | NfrA2 | 3.16 ± 1.46 | NADPH-FMN oxidoreductase | - | - |
| O32224 | AzoR2 | 3.06 ± 0.53 | similar to NAD(P)H dehydrogenase | SigA, SigG, MhqR | oxidative stress |
| P54375 | SodA | 3.00 ± 0.59 | superoxide dismutase | SigB | oxidative stress |
| P14802 | YoxD | 2.95 ± 0.61 | similar to 3-oxoacyl- acyl-carrier protein reductase | - | cell envelope |
| P80240 | GreA | 2.82 ± 0.67 | transcription elongation factor | - | transcription |
| P80864 | Tpx | 2.80 ± 0.57 | probable thiol peroxidase | Spx | oxidative stress |
| P21471 | RpsJ | 2.60 ± 0.75 | ribosomal protein uS10 | stringent response, SigA | translation |
| P39586 | YwbC | 2.55 ± 0.24 | glyoxalase I | - | oxidative stress |
| Q07836 | WapI | 2.50 ± 0.25 | immunity protein | SigA, YvrHb, WalR, DegU | - |
| P54464 | YqeY | 2.38 ± 0.24 | unknown | - | - |

**Tab. S6** MS-based protein identification for mefloquine. Protein names were taken from UniProt database [2]. Proteins were identified using the ProteinLynx Global Server software (Waters). Only proteins identified with a certainty of ≥ 95% were considered.

| **UniProt identifier** | **Name** | **MW [Da]** | **pI** | **PLGS Score** | **Number of matching peptides** | **Sequence coverage [%]** |
|---|---|---|---|---|---|---|
| O32201 | LiaH | 25698.22 | 6.18 | 14754.65 | 43 | 86.67 |
| P04990 | ThrC | 37749.08 | 5.19 | 4627.47 | 27 | 87.78 |
| O32078 | YuaE | 19110.91 | 6.20 | 6332.31 | 20 | 85.80 |
| O05394 | MccB | 40885.68 | 5.21 | 4436.64 | 21 | 51.98 |
| O07609 | YhfK | 22816.84 | 5.18 | 14905.57 | 19 | 93.46 |
| P94388 | Cah | 35806.67 | 5.42 | 6629.93 | 27 | 83.33 |
| P54569 | YqkF | 34774.41 | 5.10 | 2921.69 | 20 | 63.07 |
| P42923 | RplJ (RL10) | 18078.80 | 5.20 | 52109.85 | 26 | 94.58 |
| P54464 | YqeY | 16766.48 | 5.50 | 7717.51 | 10 | 63.51 |
| P26907 | GsiB | 13797.64 | 5.10 | 9281.11 | 9 | 52.85 |
| P80880 | TrxB | 34633.30 | 4.99 | 12393.85 | 30 | 77.22 |
| P14802 | YoxD | 25299.03 | 5.34 | 5841.61 | 19 | 78.57 |
| P81100 | YceC | 21994.88 | 5.31 | 2508.64 | 23 | 91.46 |
| P54617 | PspA | 25198.76 | 5.78 | 3056.86 | 19 | 72.69 |
| P21883 | PdhC | 35474.48 | 4.50 | 1650.19 | 14 | 47.08 |
| O34689 | MhqA | 35386.95 | 5.40 | 536.70 | 10 | 28.48 |
| P35137 | PpiB | 15370.11 | 5.50 | 4417.79 | 7 | 76.92 |
| P28368 | YvyD | 21979.76 | 5.18 | 2586.69 | 15 | 70.90 |
| P42976 | DapB | 29544.83 | 5.10 | 990.519 | 14 | 60.30 |
| P54375 | SodA | 22489.97 | 5.10 | 28005.40 | 21 | 100.00 |
| O05252 | NupN | 38418.30 | 5.10 | 3491.05 | 35 | 89.42 |
| P80864 | Tpx | 18329.77 | 4.70 | 30943.31 | 21 | 87.43 |
| P54547 | G6pD | 55803.10 | 5.28 | 3930.39 | 30 | 59.92 |
| O32029 | YrrT | 24196.28 | 5.00 | 1075.48 | 13 | 43.66 |
| P39645 | ChdC | 29562.51 | 5.00 | 2453.82 | 16 | 67.72 |
| O34305 | YtoQ | 16791.33 | 5.77 | 3606.61 | 15 | 68.92 |
| O31669 | MtnD | 20824.06 | 4.40 | 6330.56 | 10 | 46.63 |
| O06724 | YisK | 33317.12 | 5.40 | 415.65 | 14 | 63.46 |
| Q45499 | AmpS | 45798.25 | 5.00 | 9098.25 | 45 | 81.22 |
| P80244 | ClpP | 21739.07 | 5.01 | 9928.01 | 26 | 81.22 |
| P80240 | GreA | 17271.54 | 4.50 | 9430.90 | 18 | 85.35 |
| Q06756 | IspF | 17239.80 | 5.40 | 2106.45 | 6 | 34.18 |

**Tab. S7** MS-based protein identification for **4** (**BLK278**). Protein names were taken from UniProt database [2]. Proteins were identified using the ProteinLynx Global Server software (Waters). Only proteins identified with a certainty of ≥ 95% were considered.

| **UniProt identifier** | **Name** | **MW [Da]** | **pI** | **PLGS Score** | **Number of matching peptides** | **Sequence coverage [%]** |
|---|---|---|---|---|---|---|
| **O32201** | LiaH | 25698.22 | 6.18 | 13968.64 | 38 | 86.22 |
| **P81100** | YceC | 21994.88 | 5.31 | 1167.62 | 10 | 33.17 |
| **P54524** | YqiG | 40919.41 | 5.18 | 1395.06 | 19 | 55.11 |
| **P94400** | YciC | 45762.75 | 4.40 | 6935.07 | 24 | 62.72 |
| **P54617** | PspA | 25198.76 | 5.78 | 591.20 | 6 | 38.77 |
| **O05394** | MccB | 40885.67 | 5.21 | 1627.10 | 15 | 43.27 |
| **O34334** | YjoA | 17850.17 | 5.81 | 1909.94 | 6 | 42.21 |
| **O05239** | YugJ | 42903.79 | 5.22 | 5429.16 | 37 | 81.40 |
| **P46336** | IolS | 35167.94 | 5.38 | 5384.92 | 23 | 70.32 |
| **O07609** | YhfK | 22816.83 | 5.18 | 1508.23 | 9 | 55.14 |
| **P14802** | YoxD | 25299.02 | 5.34 | 1112.43 | 13 | 66.81 |
| **P32727** | NusA | 41782.32 | 4.58 | 6681.21 | 28 | 75.20 |
| **P80880** | TrxB | 34633.30 | 4.99 | 3755.39 | 19 | 68.99 |
| **P28598** | GroEL | 57424.71 | 4.53 | 240.08 | 14 | 29.23 |
| **O34833** | YceH | 41673.03 | 5.75 | 2531.11 | 34 | 81.54 |
| **P28368** | YvyD | 21979.76 | 5.18 | 336.62 | 5 | 34.39 |
| **P37573** | DisA | 40791.40 | 5.57 | 258.19 | 11 | 45.56 |
| **P80244** | ClpP | 21739.07 | 5.01 | 2683.45 | 16 | 65.99 |
| **O34305** | YtoQ | 16791.33 | 5.77 | 1271.22 | 5 | 32.43 |
| **O07603** | YhfE | 38907.05 | 5.88 | 822.76 | 12 | 34.68 |
| **P80874** | YhdN | 37369.35 | 4.77 | 1688.68 | 14 | 48.64 |

**Tab. S8** MS-based protein identification for **5** (**BLK280**). Protein names were taken from UniProt database [2]. Proteins were identified using the ProteinLynx Global Server software (Waters). Only proteins identified with a certainty of ≥ 95% were considered.

| **UniProt identifier** | **Name** | **MW [Da]** | **pI** | **PLGS Score** | **Number of matching peptides** | **Sequence coverage [%]** |
|---|---|---|---|---|---|---|
| **O32201** | LiaH | 25698.22 | 6.18 | 7517.72 | 29 | 83.11 |
| **P54617** | PspA | 25198.76 | 5.78 | 216.63 | 4 | 18.06 |
| **P39605** | NfrA1 | 28433.99 | 5.73 | 817.13 | 6 | 34.94 |
| **P54524** | YqiG | 40919.42 | 5.18 | 229.41 | 10 | 26.61 |
| **P39645** | ChdC | 29562.51 | 5.00 | 702.73 | 9 | 33.07 |
| **P52998** | PanC | 32015.42 | 4.63 | 43.94 | 5 | 20.98 |
| **P28368** | YvyD | 21979.76 | 5.18 | 565.27 | 8 | 41.27 |
| **O05394** | MccB | 40885.68 | 5.21 | 2155.74 | 16 | 46.44 |
| **O05239** | YugJ | 42903.80 | 5.22 | 1419.40 | 25 | 68.73 |
| **P80880** | TrxB | 34633.30 | 4.99 | 5260.68 | 26 | 76.27 |
| **P46336** | IolS | 35167.95 | 5.38 | 1286.73 | 18 | 65.81 |
| **O34334** | YjoA | 17850.17 | 5.81 | 3387.21 | 9 | 59.74 |
| **P80871** | YwrO | 20011.76 | 5.20 | 2364.19 | 10 | 60.00 |
| **O30509** | GatB | 53724.29 | 4.90 | 2568.39 | 29 | 59.24 |
| **P81100** | YceC | 21994.88 | 5.31 | 2276.56 | 14 | 60.30 |
| **O34833** | YceH | 41673.03 | 5.75 | 1411.66 | 26 | 63.91 |
| **P37967** | PnbA | 54100.27 | 4.78 | 1286.61 | 13 | 35.38 |
| **O07609** | YhfK | 22816.84 | 5.18 | 2938.95 | 13 | 78.50 |
| **P14802** | YoxD | 25299.03 | 5.34 | 1457.65 | 15 | 69.75 |
| **P80244** | ClpP | 21739.07 | 5.01 | 506.55 | 6 | 26.90 |
| **P80874** | YhdN | 37369.36 | 4.77 | 578.96 | 13 | 42.30 |
| **P28598** | GroEL | 57424.71 | 4.53 | 6906.52 | 51 | 79.04 |

**Tab. S9** MS-based protein identification for **6** (**TKK088**). Protein names were taken from UniProt database [2]. Proteins were identified using the ProteinLynx Global Server software (Waters). Only proteins identified with a certainty of ≥ 95% were considered.

| **UniProt identifier** | **Name** | **MW [Da]** | **pI** | **PLGS Score** | **Number of matching peptides** | **Sequence coverage [%]** |
|---|---|---|---|---|---|---|
| O32201 | LiaH | 25698.22 | 6.18 | 31279.69 | 50 | 91.11 |
| P54617 | PspA | 25198.76 | 5.78 | 4557.40 | 22 | 68.72 |
| P31847 | YpuA | 31294.60 | 4.50 | 5511.73 | 14 | 54.48 |
| O31711 | YknY | 25385.95 | 5.80 | 1816.69 | 12 | 55.65 |
| P81100 | YceC | 21994.88 | 5.31 | 6253.16 | 29 | 92.46 |
| P02394 | RplJ (RL7) | 12750.67 | 4.40 | 99999.80 | 24 | 98.37 |
| P04990 | ThrC | 37749.08 | 5.19 | 19538.01 | 38 | 95.74 |
| O32078 | YuaE | 19110.91 | 6.20 | 6157.08 | 20 | 84.57 |
| O05239 | YugJ | 42903.80 | 5.22 | 13316.40 | 42 | 87.86 |
| P39605 | NfrA1 | 28433.99 | 5.73 | 902.30 | 14 | 64.26 |
| P14802 | YoxD | 25299.03 | 5.34 | 13860.16 | 21 | 80.25 |
| P54569 | YqkF | 34774.41 | 5.10 | 5072.93 | 31 | 79.74 |
| P28368 | YvyD | 21979.76 | 5.18 | 947.34 | 14 | 51.32 |
| P39645 | ChdC | 29562.51 | 5.00 | 1760.35 | 18 | 66.54 |
| P42923 | RplJ (RL10) | 18078.80 | 5.20 | 50794.79 | 25 | 94.58 |
| O34969 | YfjR | 30531.16 | 4.90 | 2006.93 | 16 | 55.59 |
| P80244 | ClpP | 21739.07 | 5.01 | 10590.71 | 24 | 80.20 |
| P38493 | Cmk | 25096.52 | 5.00 | 4429.86 | 28 | 87.50 |
| P23446 | FlgG | 27455.41 | 4.70 | 1004.45 | 16 | 83.33 |
| O07607 | YhfI | 26753.00 | 5.60 | 748.34 | 7 | 33.20 |
| O31618 | ThiG | 27136.34 | 4.70 | 10671.02 | 27 | 79.69 |
| P54464 | YqeY | 16766.48 | 5.50 | 11831.11 | 17 | 85.14 |
| O32224 | AzoR2 | 23272.27 | 5.10 | 6746.58 | 16 | 72.04 |
| P32396 | HemH | 35404.90 | 4.60 | 253.656 | 9 | 31.61 |
| P37573 | DisA | 40791.41 | 5.60 | 169.20 | 13 | 29.17 |
| P80864 | Tpx | 18329.77 | 4.70 | 18343.75 | 16 | 87.43 |
| P54375 | SodA | 22489.97 | 5.10 | 29529.34 | 17 | 82.67 |
| P80240 | GreA | 17271.54 | 4.50 | 6858.48 | 17 | 85.99 |
| P94424 | NfrA2 | 27924.60 | 5.10 | 1418.75 | 6 | 36.14 |
| O05394 | MccB | 40885.68 | 5.21 | 400.10 | 10 | 25.59 |

**Tab. S10** MS-based protein identification for **7** (**TKK129**). Protein names were taken from UniProt database [2]. Proteins were identified using the ProteinLynx Global Server software (Waters). Only proteins identified with a certainty of ≥ 95% were considered.

| UniProt identifier | Name | MW [Da] | pI | PLGS Score | Number of matching peptides | Sequence coverage [%] |
|---|---|---|---|---|---|---|
| O32201 | LiaH | 25698.22 | 6.18 | 30172.17 | 48 | 92.44 |
| P71088 | Spo0M | 29790.43 | 4.30 | 1005.40 | 13 | 48.84 |
| P54617 | PspA | 25198.76 | 5.78 | 3216.77 | 34 | 82.82 |
| P31847 | YpuA | 31294.60 | 4.50 | 7678.14 | 26 | 74.83 |
| P02394 | RplJ (RL7) | 12750.67 | 4.40 | 106265.90 | 21 | 97.56 |
| O05239 | YugJ | 42903.80 | 5.22 | 8703.370 | 48 | 97.16 |
| P39605 | NfrA1 | 28433.99 | 5.73 | 6025.48 | 22 | 70.28 |
| P04990 | ThrC | 37749.08 | 5.19 | 28106.68 | 40 | 96.31 |
| P54569 | YqkF | 34774.41 | 5.10 | 4470.06 | 27 | 69.93 |
| P81100 | YceC | 21994.88 | 5.31 | 4083.63 | 26 | 75.88 |
| O32078 | YuaE | 19110.91 | 6.20 | 11377.46 | 19 | 82.10 |
| P39645 | ChdC | 29562.51 | 5.00 | 3515.67 | 23 | 87.01 |
| P42923 | RplJ (RL10) | 18078.80 | 5.20 | 45666.54 | 24 | 94.58 |
| P49786 | AccB | 17285.61 | 4.40 | 3770.26 | 14 | 92.45 |
| O05240 | YugK | 43577.49 | 4.60 | 376.23 | 15 | 40.00 |
| P94424 | NfrA2 | 27924.60 | 5.10 | 1686.65 | 9 | 46.18 |
| O32224 | AzoR2 | 23272.27 | 5.10 | 16671.22 | 24 | 91.94 |
| P54375 | SodA | 22489.97 | 5.10 | 38944.89 | 23 | 100.00 |
| P14802 | YoxD | 25299.03 | 5.34 | 9140.26 | 31 | 92.02 |
| P80240 | GreA | 17271.54 | 4.50 | 12339.35 | 19 | 87.26 |
| P80864 | Tpx | 18329.77 | 4.70 | 33883.55 | 20 | 92.81 |
| P21471 | RpsJ | 11665.65 | 10.20 | 9375.58 | 16 | 75.49 |
| P39586 | YwbC | 14433.35 | 4.43 | 1851.85 | 5 | 80.95 |
| Q07836 | WapI | 16542.85 | 4.40 | 5443.79 | 15 | 80.99 |
| P54464 | YqeY | 16766.48 | 5.50 | 19138.15 | 21 | 85.81 |

#### Supplementary Figures

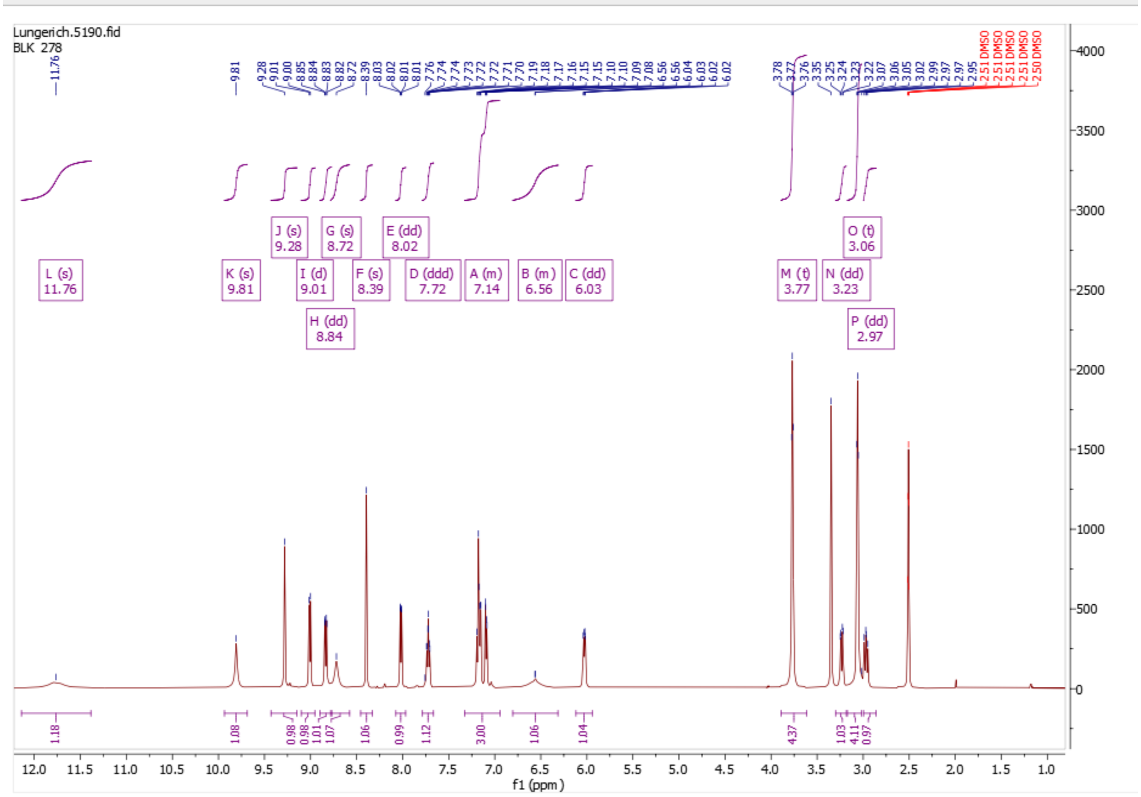


**Supplementary Fig. S1**: $^{1}$H NMR spectrum (600 MHz, DMSO-d6) of **4** (**BLK278**).

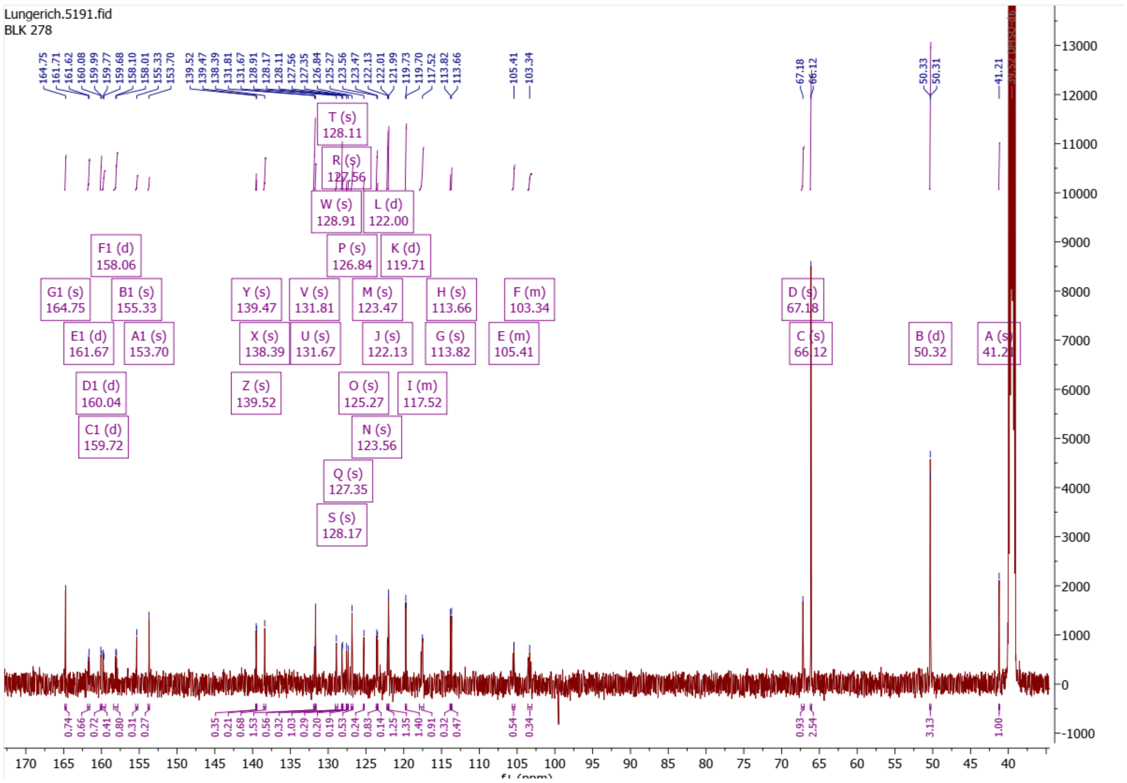

**Supplementary Fig. S2**: $^{13}C$ NMR spectrum (151 MHz, DMSO-d6) of **4** (**BLK278**).

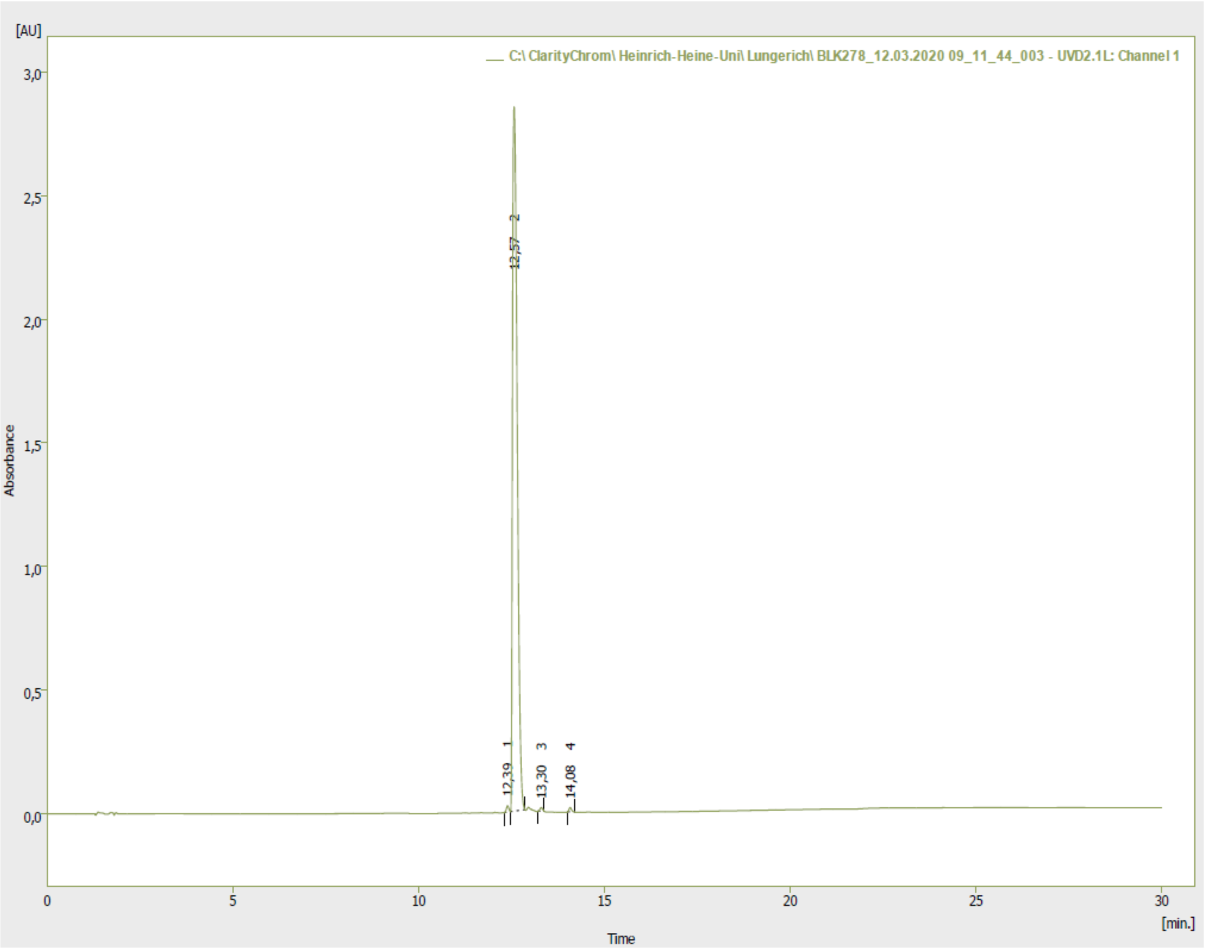


**Supplementary Fig. S3**: HPLC chromatogram of the purity analysis of **4** (**BLK278**).

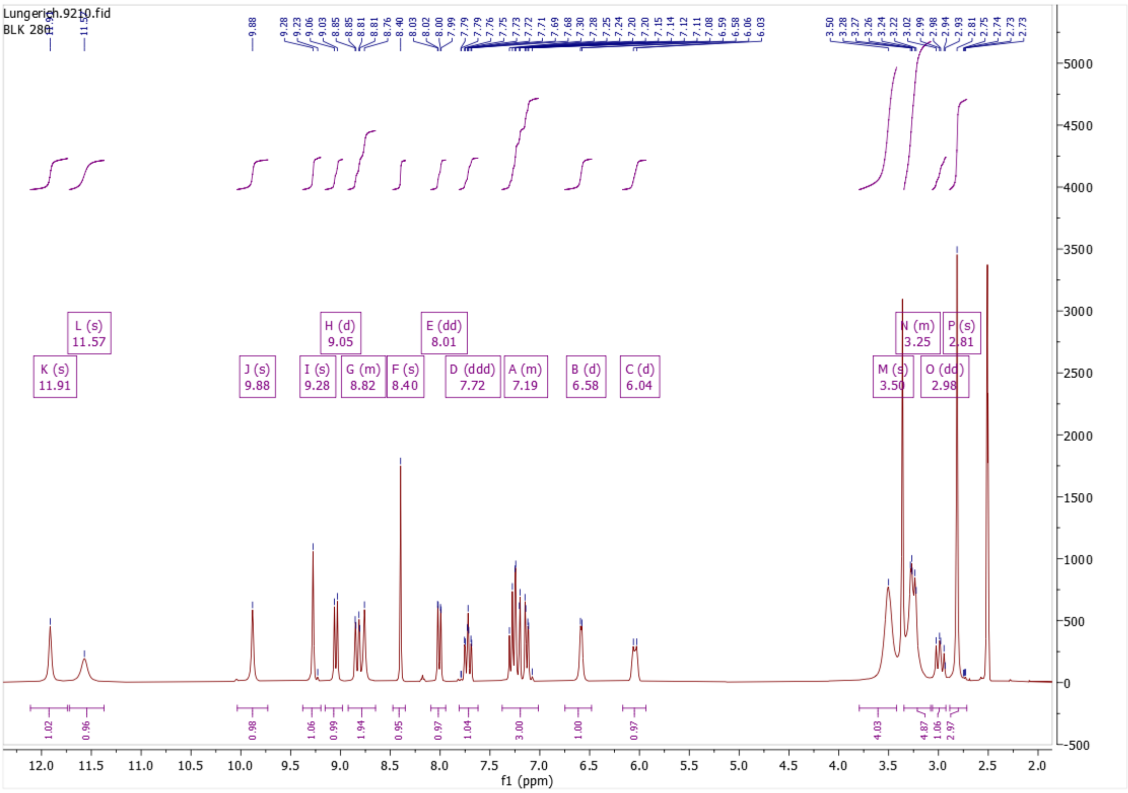


**Supplementary Fig. S4**: $^{1}$H NMR spectrum (600 MHz, DMSO-d6) of **5** (**BLK280**).

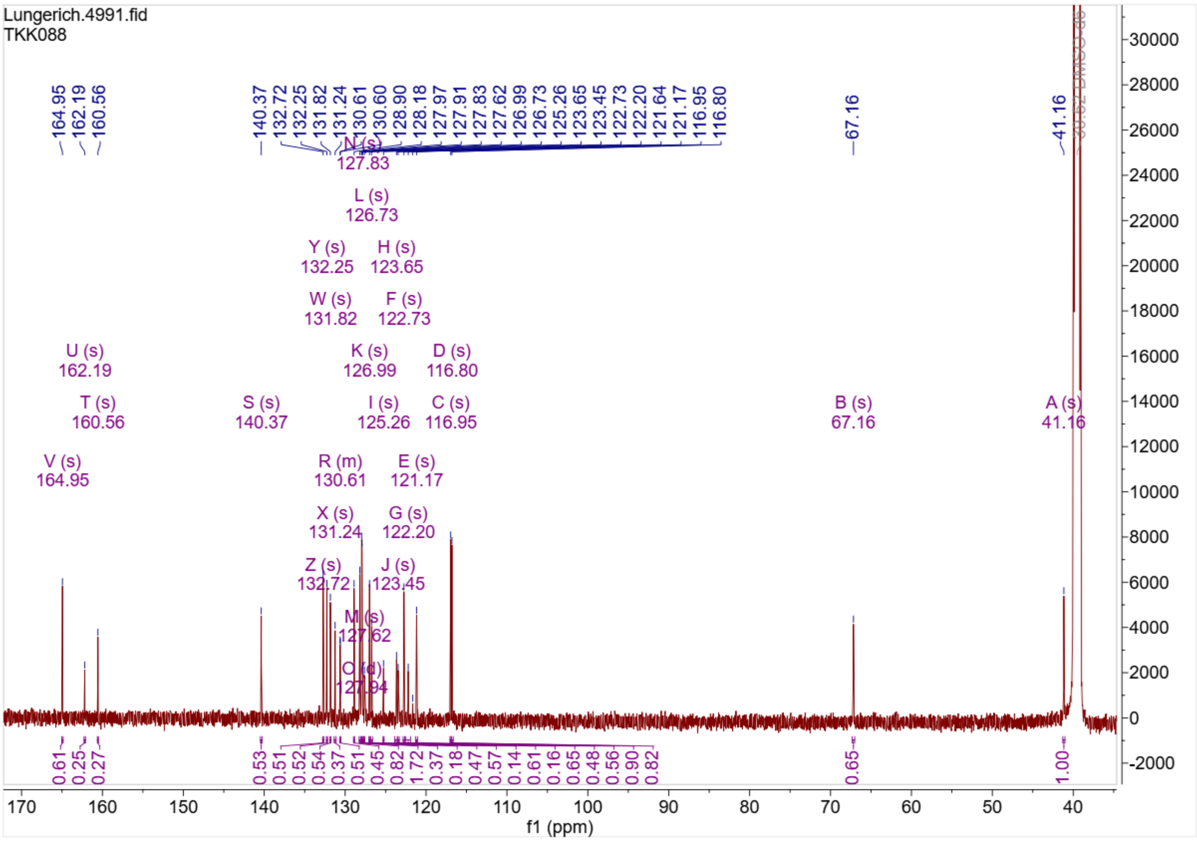


**Supplementary Fig. S5**: $^{13}$C NMR spectrum (151 MHz, DMSO-d6) of **5** (**BLK280**).

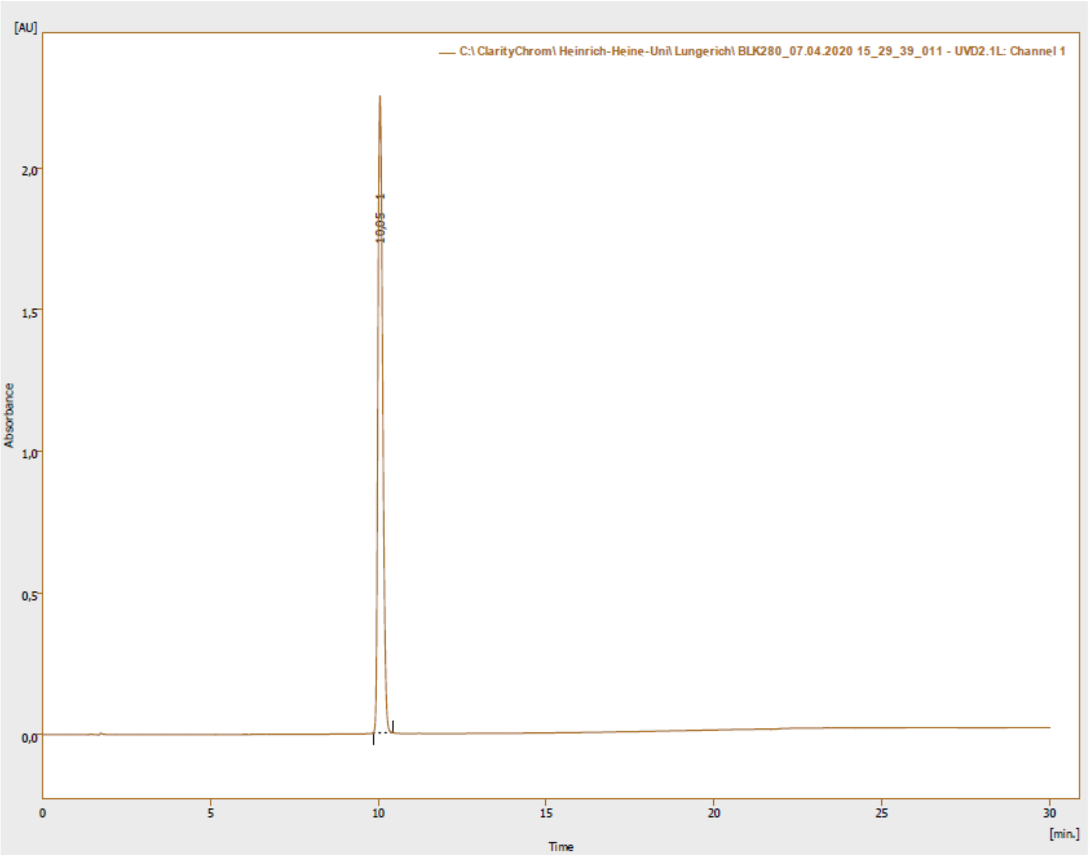


**Supplementary Fig. S6**: HPLC chromatogram of the purity analysis of **5** (**BLK280**).

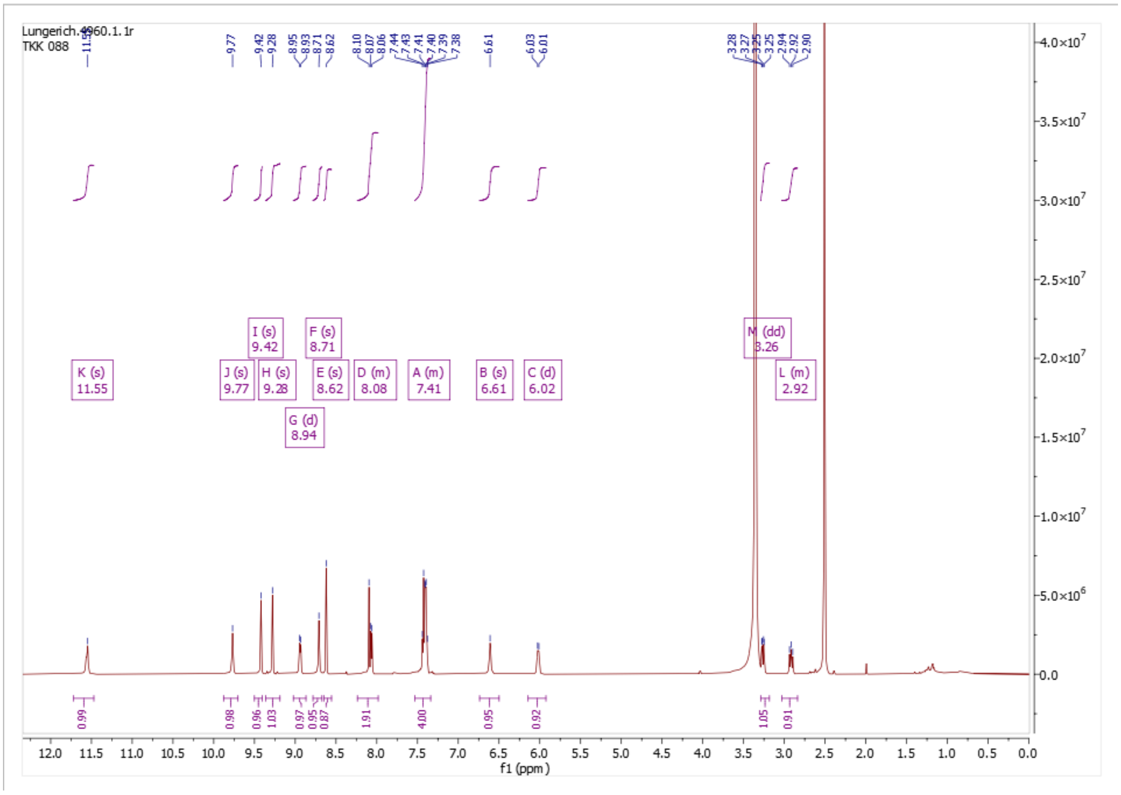


**Supplementary Fig. S7**: $^{1}$H NMR spectrum (600 MHz, DMSO-d6) of **6** (**TKK088**).

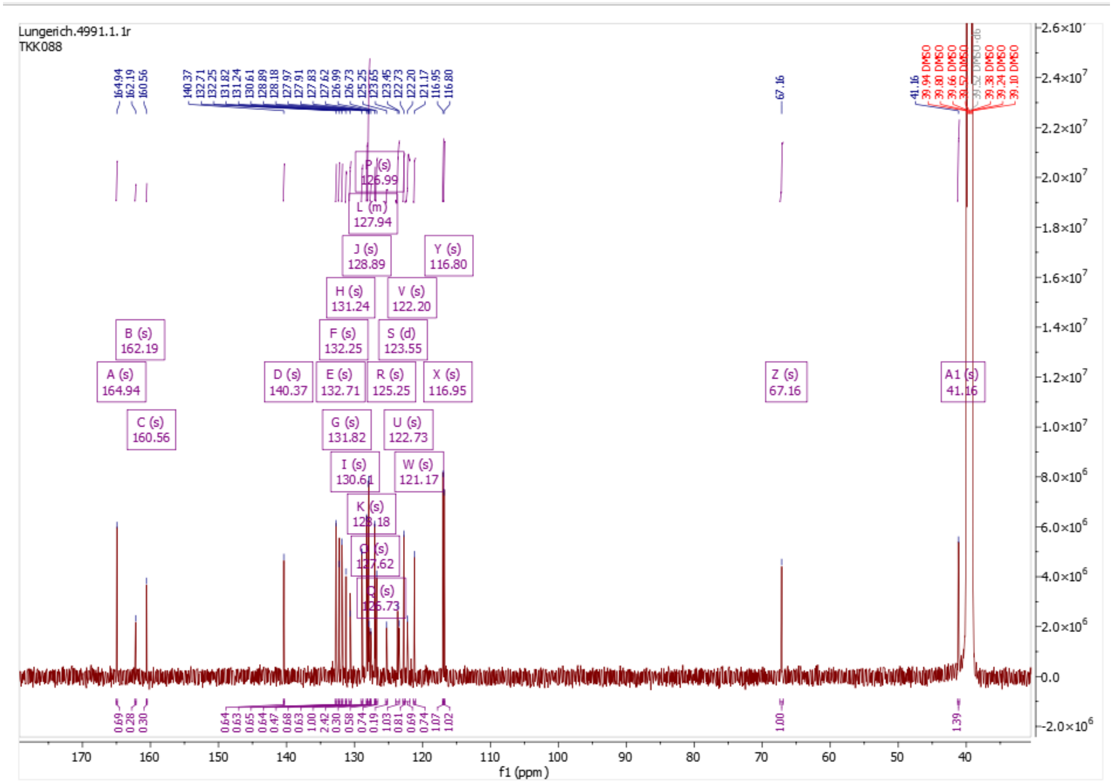


**Supplementary Fig. S8**: $^{13}$C NMR spectrum (151 MHz, DMSO-d6) of **6** (**TKK088**).

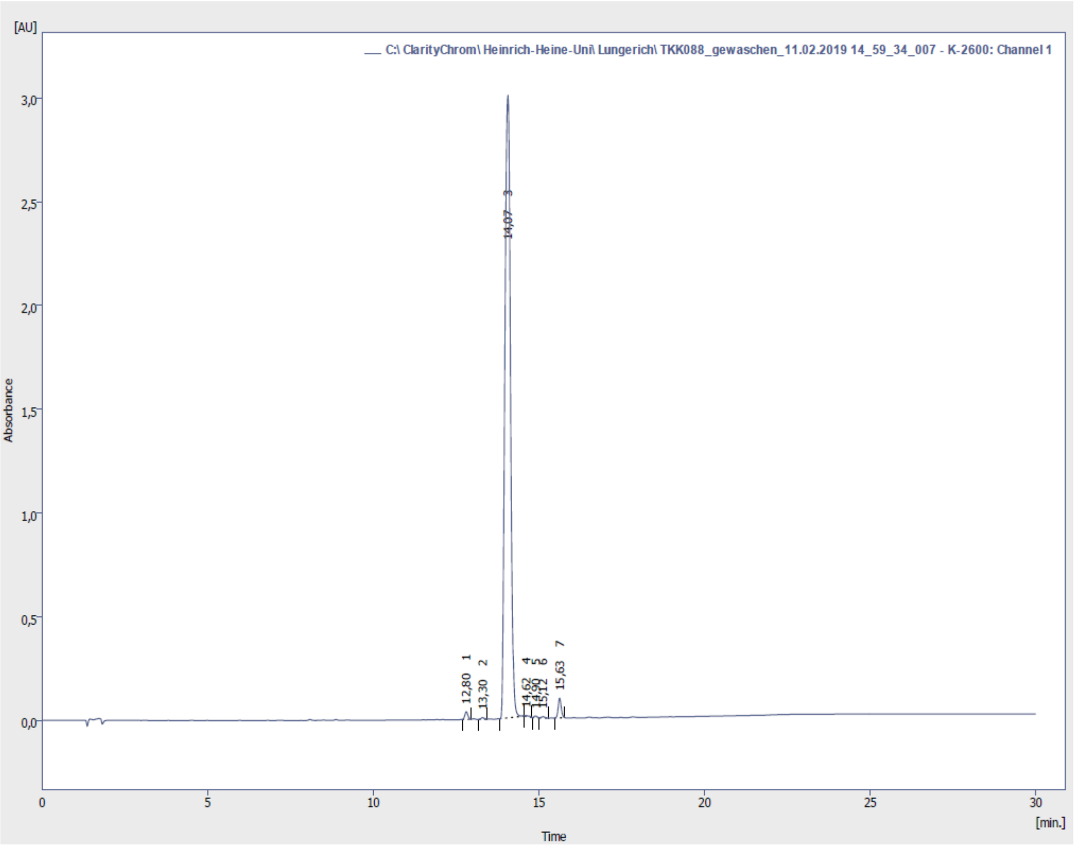


**Supplementary Fig. S9**: HPLC chromatogram of the purity analysis of **6** (**TKK088**).

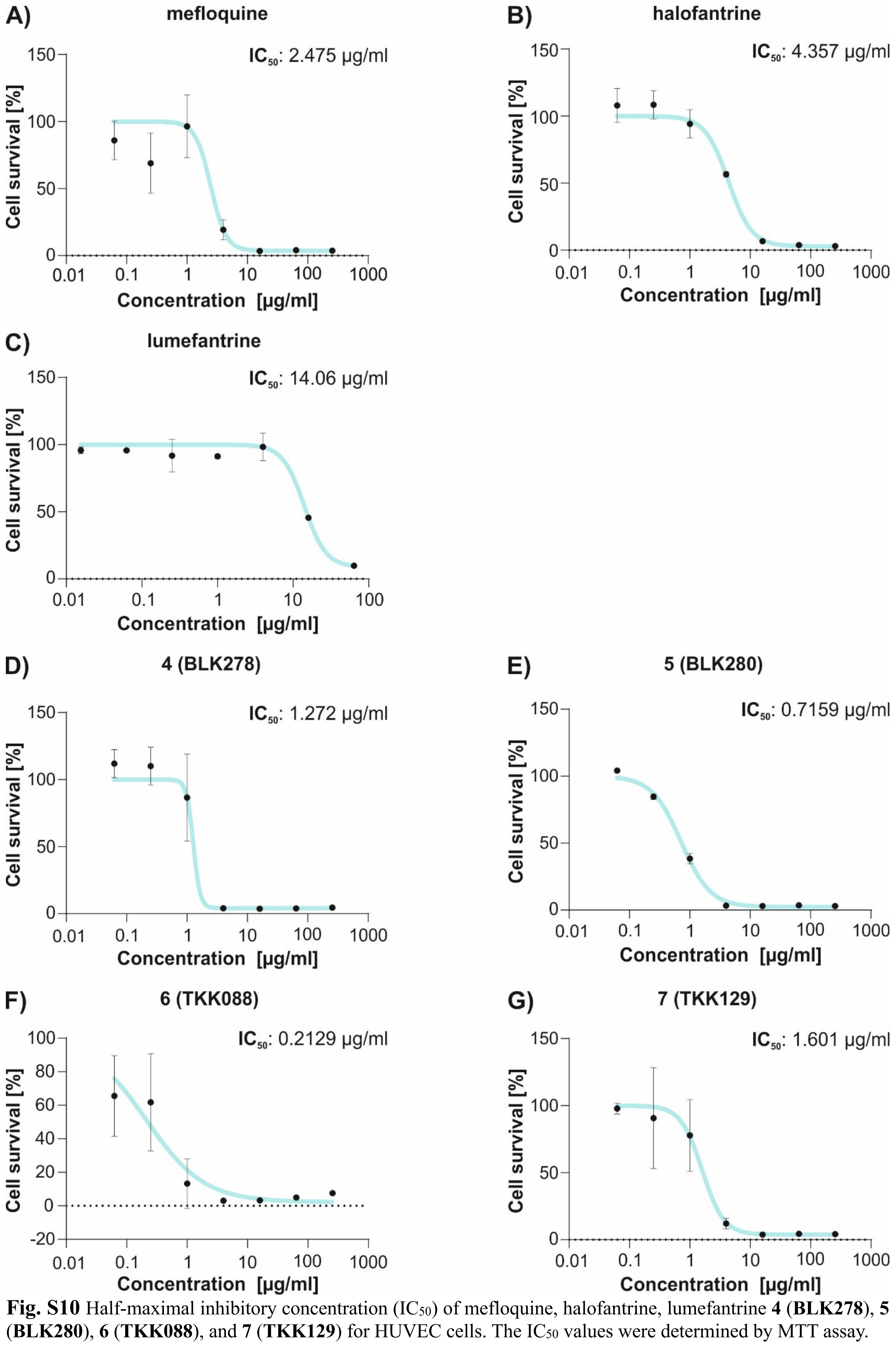


**Fig. S10** Half-maximal inhibitory concentration ($IC_{50}$) of mefloquine, halofantrine, lumefantrine **4** (**BLK278**), **5** (**BLK280**), **6** (**TKK088**), and **7** (**TKK129**) for HUVEC cells. The $IC_{50}$ values were determined by MTT assay.

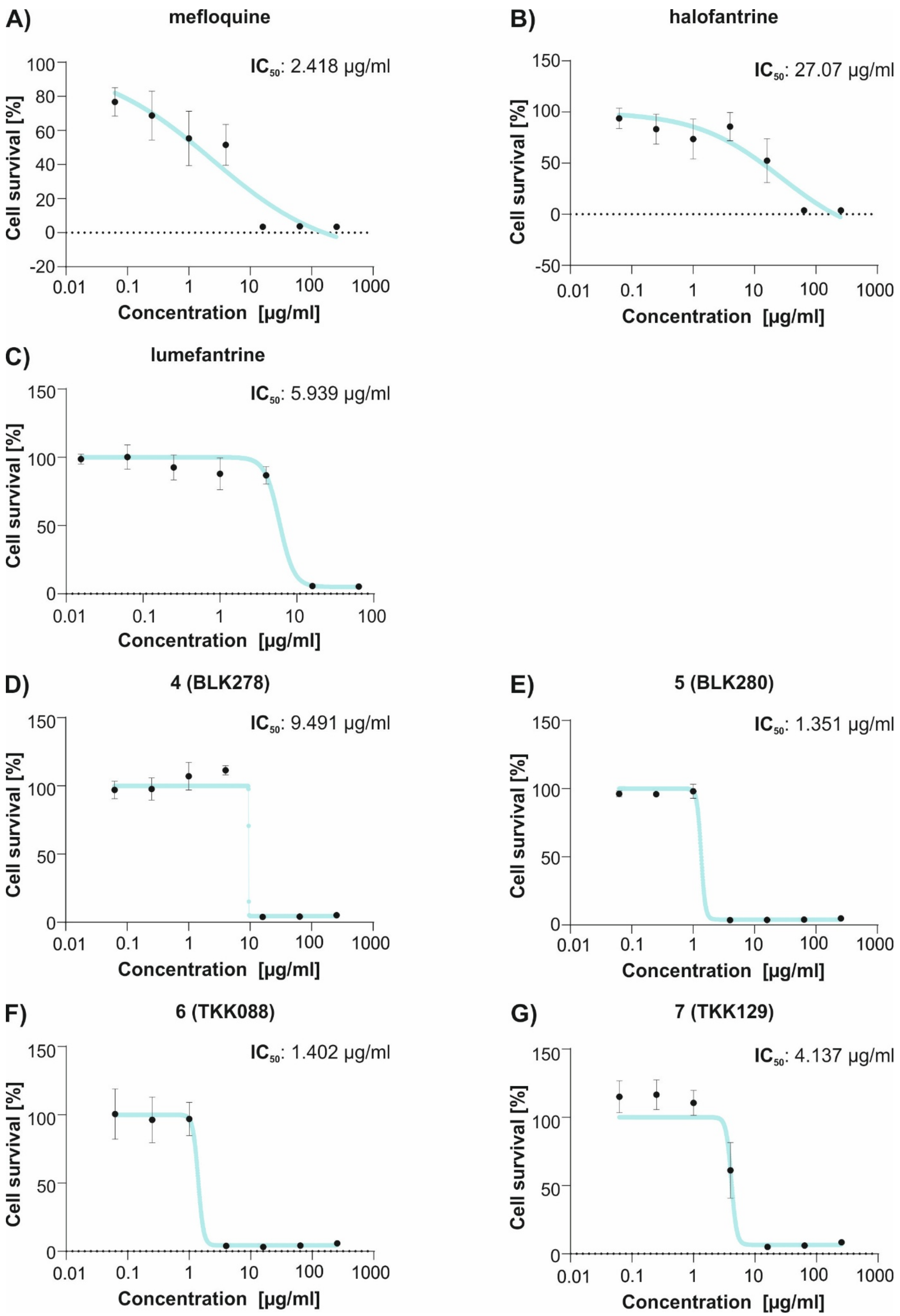


**Fig. S11** Half-maximal inhibitory concentration ($IC_{50}$) of mefloquine, halofantrine, lumefantrine **4** (**BLK278**), **5** (**BLK280**), **6** (**TKK088**), and **7** (**TKK129**) for HAOSMC cells. The $IC_{50}$ values were determined by MTT assay.

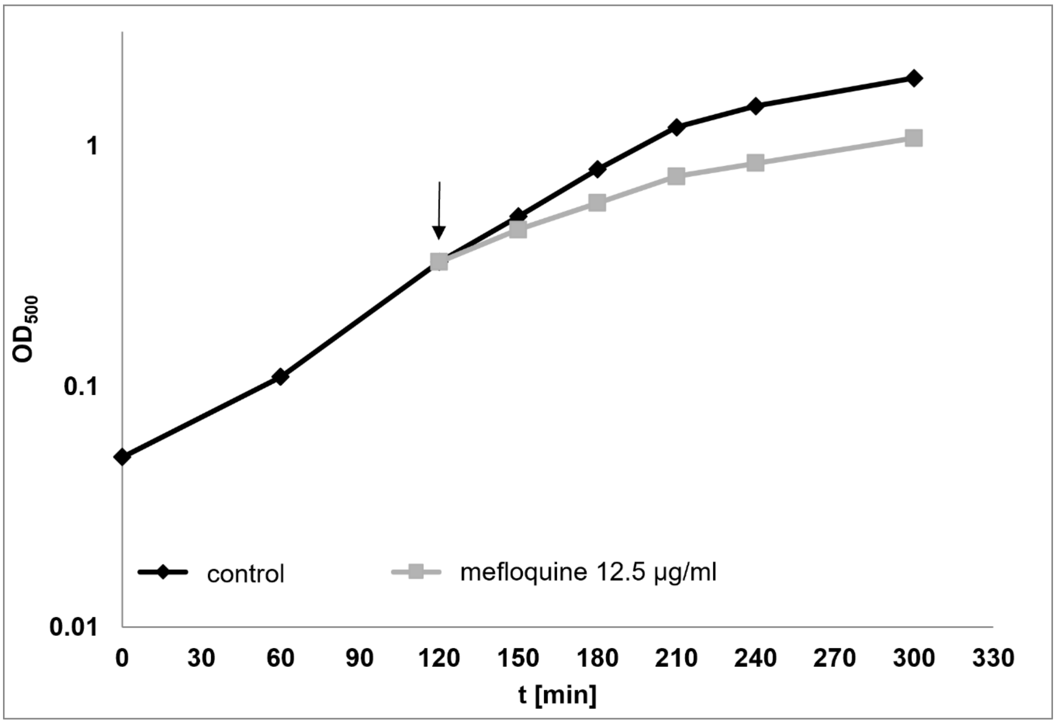


**Fig. S12** Inhibition of logarithmically growing *B. subtilis* 168 exposed to 12.5 µg/ml mefloquine. Effect of 12.5 µg/ml mefloquine on the growth of *B. subtilis* 168 in BMM minimal medium. Growth was monitored in parallel to the L-[$^{35}$S]-methioine pulse labeling, in which proteins were labeled for 5 min starting 10 min after antibiotic addition. The arrow marks the time of antibiotic addition. Data shown are representative of three independent biological replicates.

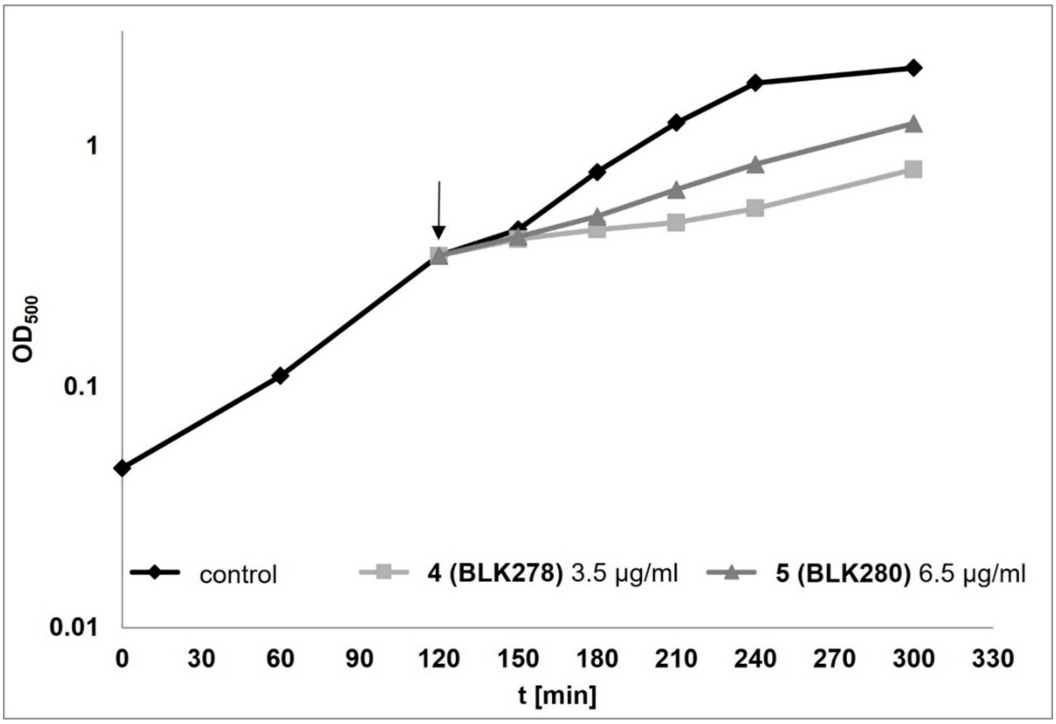


**Fig. S13** Inhibition of logarithmically growing *B. subtilis* 168 exposed to 3.5 µg/ml **4** (**BLK278**) or 6.5 µg/ml **5** (**BLK280**). Effect of 3.5 µg/ml **4** (**BLK278**) and 6.5 µg/ml **5** (**BLK280**) on the growth of *B. subtilis* 168 in BMM minimal medium. Growth was monitored in parallel to the L-[$^{35}$S]-methioine pulse labeling, in which proteins were labeled for 5 min starting 10 min after antibiotic addition. The arrow marks the time of antibiotic addition. Data shown are representative of three independent biological replicates.

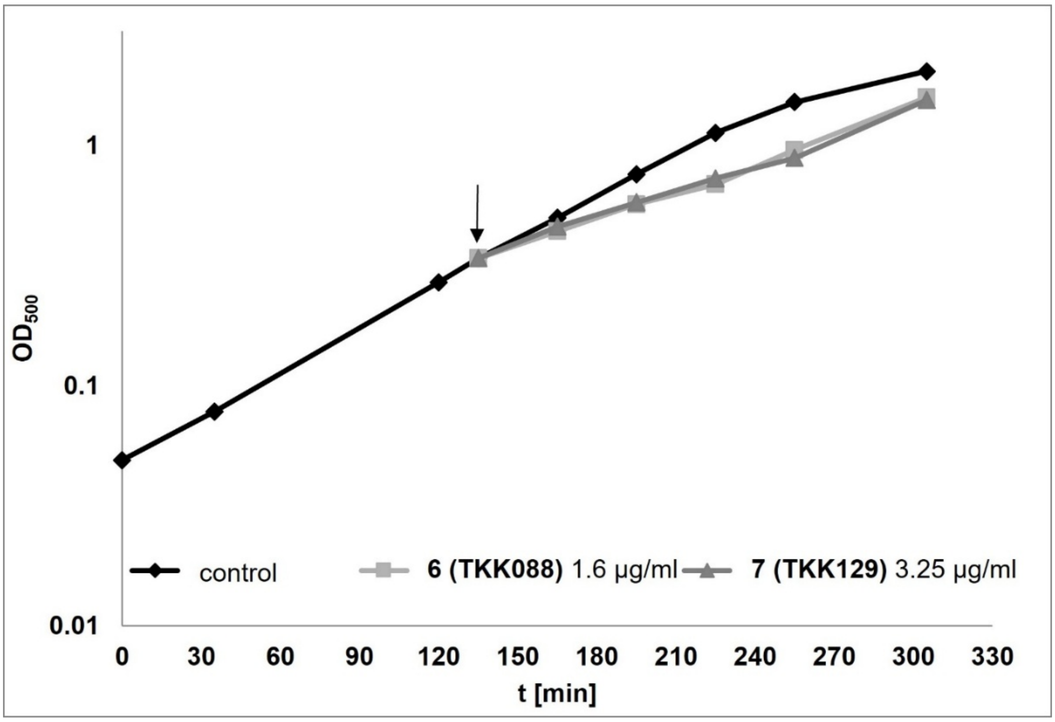


**Fig. S14** Inhibition of logarithmically growing *B. subtilis* 168 exposed to 1.6 µg/ml **6** (**TKK088**) or 3.5 µg/ml **7** (**TKK129**). Effect of 3.25 µg/ml **7** (**TKK129**) or 1.6 µg/ml **6** (**TKK088**) on the growth of *B. subtilis* 168 in BMM minimal medium. Growth was monitored in parallel to the L-[$^{35}$S]-methioine pulse labeling, in which proteins were labeled for 5 min starting 10 min after antibiotic addition. The arrow marks the time of antibiotic addition. Data shown are representative of three independent biological replicates.

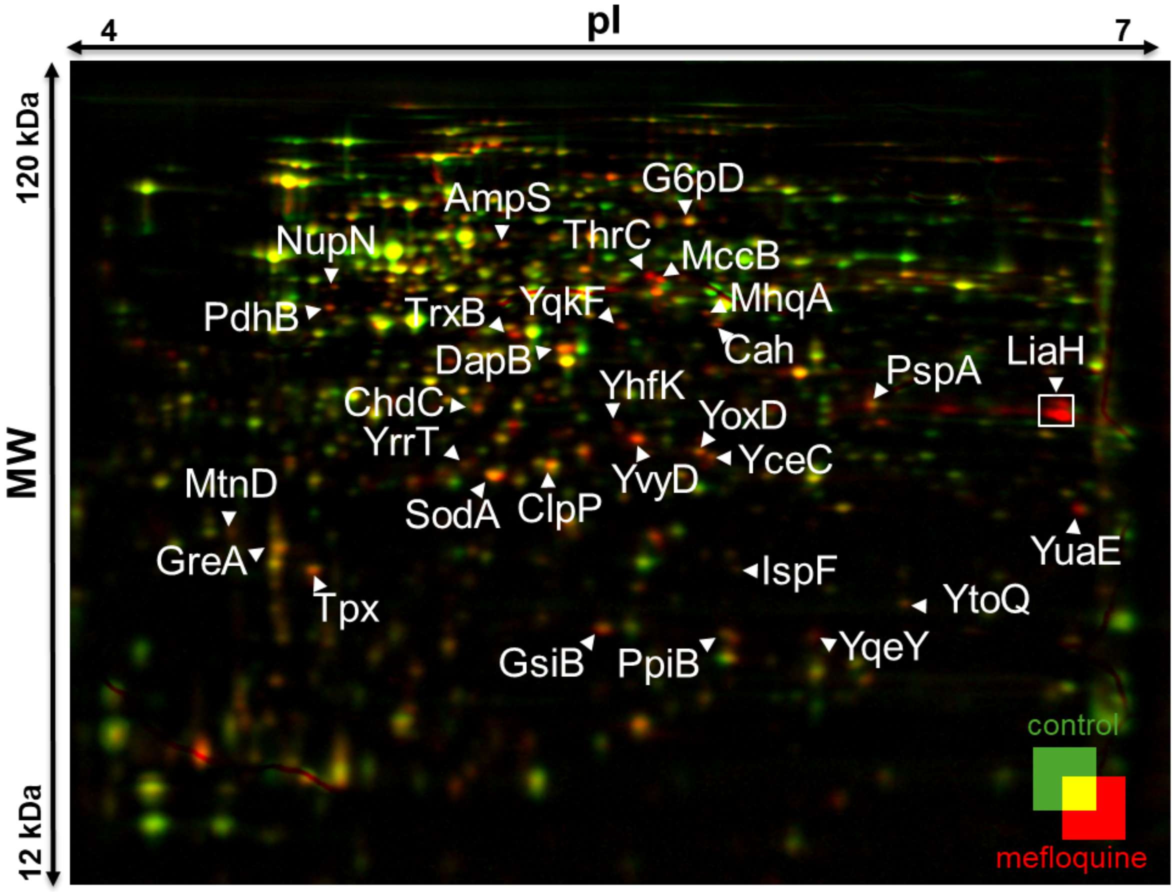


**Fig. S15** Proteomic response of *B. subtilis* 168 treated with 12.5 µg/ml mefloquine. The false-color image overlay depicts the proteomic response of *B. subtilis* 168 to 12.5 µg/ml mefloquine (red) in comparison to an untreated control (green). *B. subtilis* was treated with mefloquine for 15 min in total. After 10 min of treatment, L-[$^{35}$S]-methionine was added to treated and control cultures to pulse-label newly synthesized proteins. Labeling was terminated with chloramphenicol and an excess of L-methionine. After cell lysis, cytosolic proteins were separated first by isoelectric focusing based on the isoelectric point (pI), second by mass using SDS-PAGE. After drying the gels, the proteomic response was documented as an overlay of two autoradiographs. Spots with ≥2 fold increase in relative intensity were identified with the software Delta2D and subjected to protein identification. To this end spots cut out from a preparative gel underwent in-gel digestion and identification by LC/MS-MS. Shown are the marker proteins identified. The data shown represent three independent biological replicates.

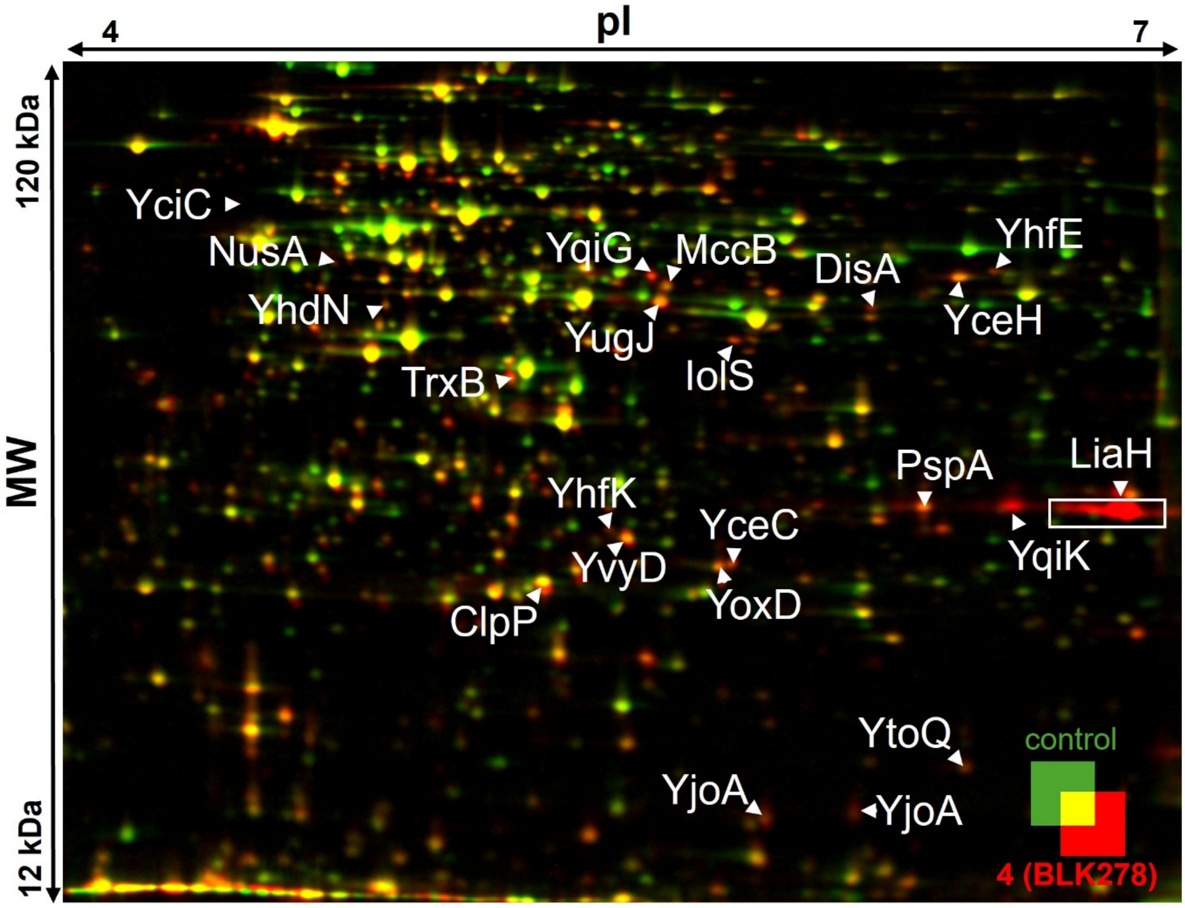


**Fig. S16** Proteomic response of *B. subtilis* 168 treated with 3.5 µg/ml **4** (**BLK278**). The false-color image overlay depicts the proteomic response of *B. subtilis* 168 to 3.5 µg/ml **4** (**BLK278**) (red) in comparison to an untreated control (green). *B. subtilis* was treated with **4** (**BLK278**) for 15 min in total. After 10 min of treatment, L-[$^{35}$S]-methionine was added to treated and control cultures to pulse-label newly synthesized proteins. Labeling was terminated with chloramphenicol and an excess of L-methionine. After cell lysis, cytosolic proteins were separated first by isoelectric focusing based on the isoelectric point (pI), second by mass using SDS-PAGE. After drying the gels, the proteomic response was documented as an overlay of two autoradiographs. Spots with ≥2 fold increase in relative intensity were identified with the software Delta2D and subjected to protein identification. To this end spots cut out from a preparative gel underwent in-gel digestion and identification by LC/MS-MS. Shown are the marker proteins identified. The data shown represent three independent biological replicates.

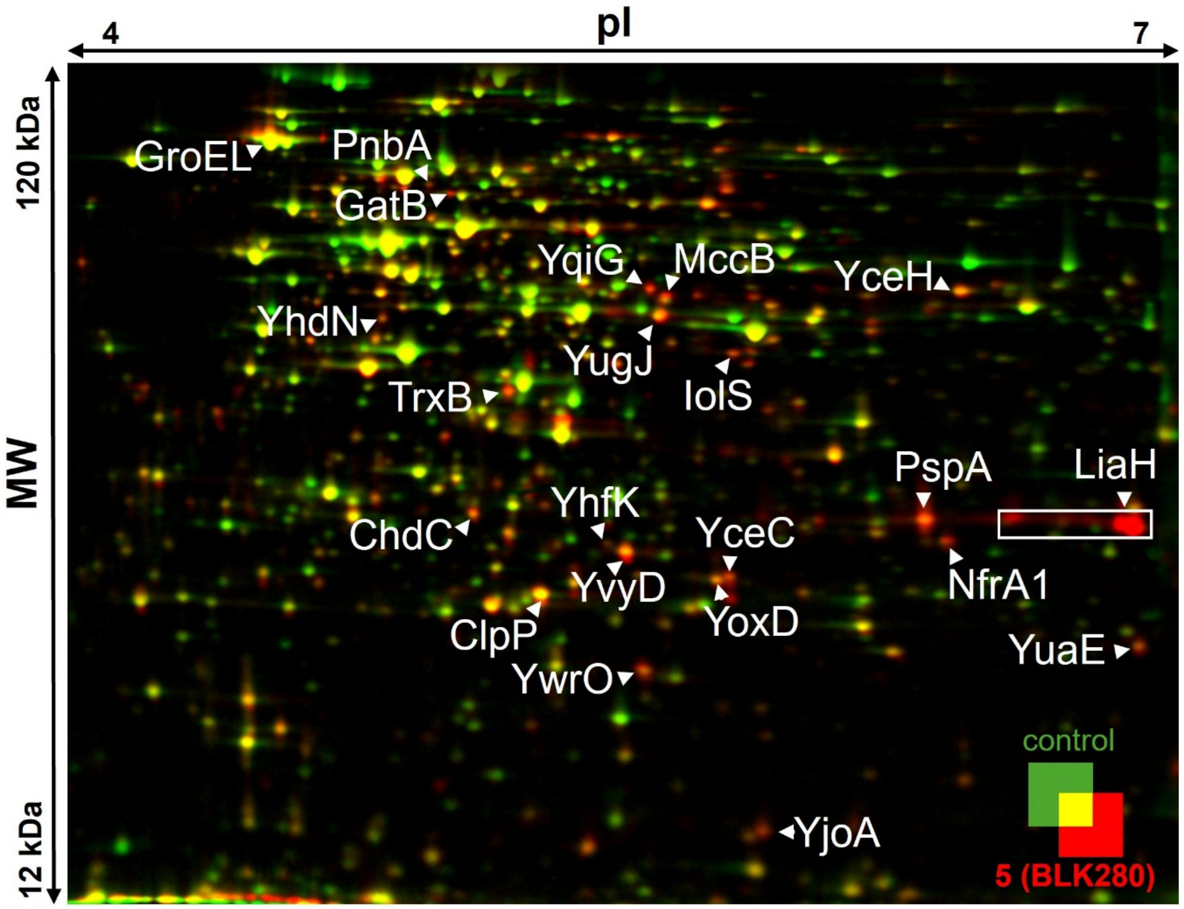


**Fig. S17** Proteomic response of *B. subtilis* 168 treated with 6.5 µg/ml **5 (BLK280)**. The false-color image overlay depicts the proteomic response of *B. subtilis* 168 to 6.5 µg/ml **5 (BLK280)** (red) in comparison to an untreated control (green). *B. subtilis* was treated with **5 (BLK280)** for 15 min in total. After 10 min of treatment, L-[$^{35}$S]-methionine was added to treated and control cultures to pulse-label newly synthesized proteins. Labeling was terminated with chloramphenicol and an excess of L-methionine. After cell lysis, cytosolic proteins were separated first by isoelectric focusing based on the isoelectric point (pI), second by mass using SDS-PAGE. After drying the gels, the proteomic response was documented as an overlay of two autoradiographs. Spots with ≥2 fold increase in relative intensity were identified with the software Delta2D and subjected to protein identification. To this end spots cut out from a preparative gel underwent in-gel digestion and identification by LC/MS-MS. Shown are the marker proteins identified. The data shown represent three independent biological replicates.

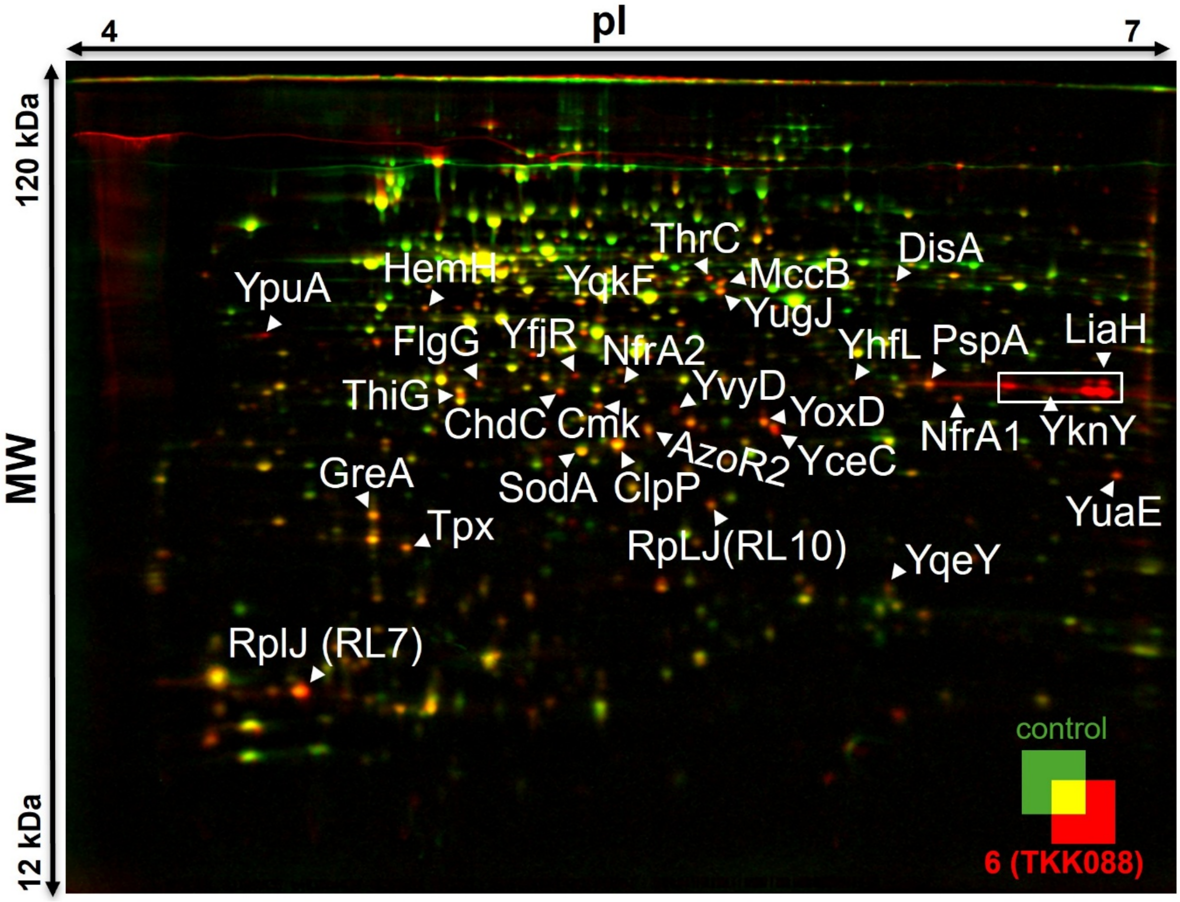


**Fig. S18** Proteomic response of *B. subtilis* 168 treated with 1.6 µg/ml **6** (**TKK088**). The false-color image overlay depicts the proteomic response of *B. subtilis* 168 to 1.6 µg/ml **6** (**TKK088**) (red) in comparison to an untreated control (green). *B. subtilis* was treated with **6** (**TKK088**) for 15 min in total. After 10 min of treatment, L-[$^{35}$S]-methionine was added to treated and control cultures to pulse-label newly synthesized proteins. Labeling was terminated with chloramphenicol and an excess of L-methionine. After cell lysis, cytosolic proteins were separated first by isoelectric focusing based on the isoelectric point (pI), second by mass using SDS-PAGE. After drying the gels, the proteomic response was documented as an overlay of two autoradiographs. Spots with ≥2 fold increase in relative intensity were identified with the software Delta2D and subjected to protein identification. To this end spots cut out from a preparative gel underwent in-gel digestion and identification by LC/MS-MS. Shown are the marker proteins identified. The data shown represent three independent biological replicates.

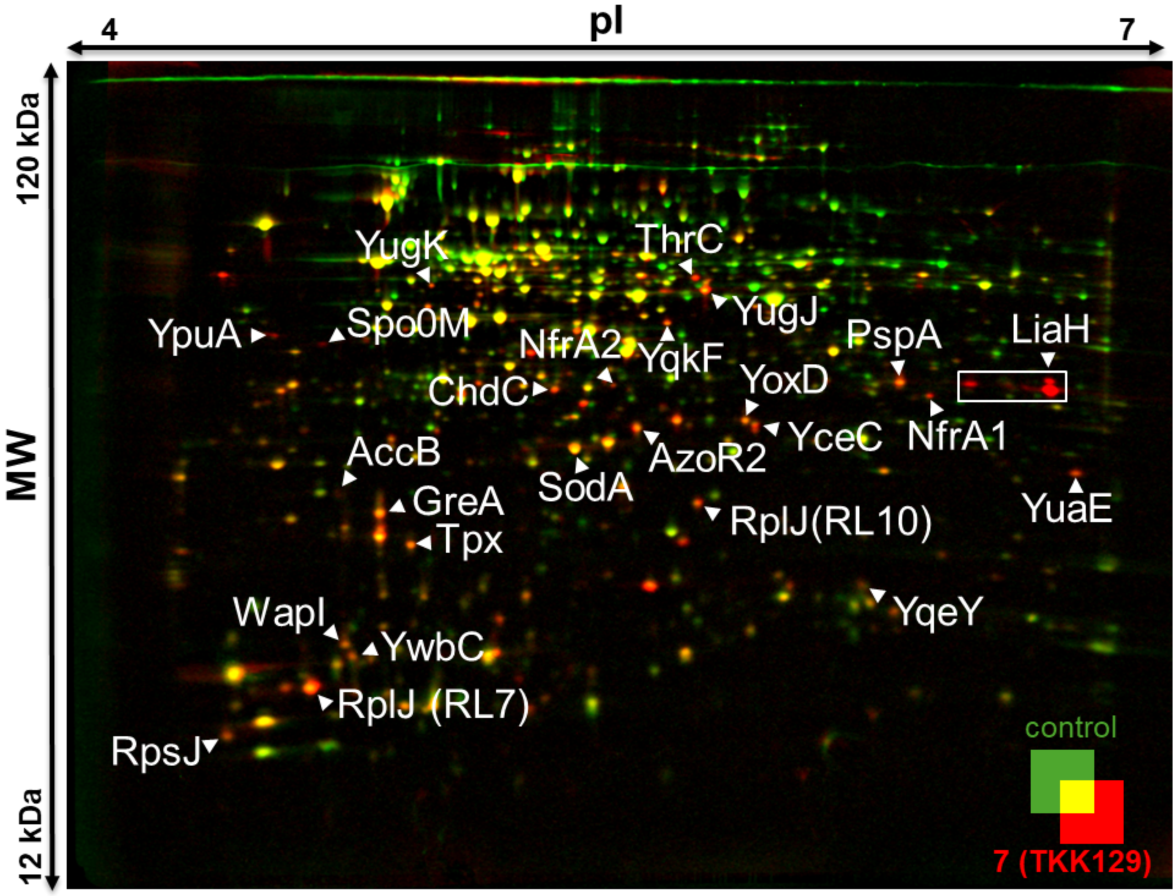


**Fig. S19** Proteomic response of *B. subtilis* 168 treated with 3.25 µg/ml **7** (**TKK129**). The false-color image overlay reflects protein synthesis of *B. subtilis* 168 after exposure to 3.25 µg/ml **7** (**TKK129**) (red) in comparison to an untreated control (green). *B. subtilis* was treated with **7** (**TKK129**) for 15 min in total. After 10 min of treatment, L-[$^{35}$S]-methionine was added to treated and control cultures to pulse-label newly synthesized proteins. Labeling was terminated with chloramphenicol and an excess of L-methionine. After cell lysis, cytosolic proteins were separated first by isoelectric focusing based on the isoelectric point (pI), second by mass using SDS-PAGE. After drying the gels, the proteomic response was documented as an overlay of two autoradiographs. Spots with ≥2 fold increase in relative intensity were identified with the software Delta2D and subjected to protein identification. To this end spots cut out from a preparative gel underwent in-gel digestion and identification by LC/MS-MS. Shown are the marker proteins identified. The data shown represent three independent biological replicates.
